# Fibroblasts Regulate the Transformation Potential of Human Papillomavirus-positive Keratinocytes

**DOI:** 10.1101/2024.09.16.613347

**Authors:** Claire D. James, Rachel L. Lewis, Austin J. Witt, Christiane Carter, Nabiha M. Rais, Xu Wang, Molly L. Bristol

**Author notes:** Address correspondence to Molly L. Bristol.

## Abstract

Persistent human papillomavirus (HPV) infection is necessary but insufficient for viral oncogenesis. Additional contributing co-factors, such as immune evasion and viral integration have been implicated in HPV-induced cancer progression. It is widely accepted that HPV+ keratinocytes require co-culture with fibroblasts to maintain viral episome expression, yet the exact mechanisms for this have yet to be elucidated. Here we present comprehensive RNA sequencing and proteomic analysis demonstrating that fibroblasts not only support the viral life cycle, but reduce HPV+ keratinocyte transformation. Our co-culture models offer novel insights into HPV-related transformation mechanisms.

**Highlights:**

- Fibroblasts support HPV RNA expression and episomal maintenance in HPV+ keratinocytes
- Fibroblasts reduce EMT related expression in HPV+ keratinocytes
- Fibroblasts promote EMT related expression in E6E7+ keratinocytes

## 1 Introduction

Human papillomaviruses (HPVs) infect the basal keratinocytes of differentiating squamous epithelia [1]. Some current estimates suggest there may be more than 400 types of HPV, however, there are approximately 12 high-risk HPV types with the capacity to cause cancer in the general population [2–4]. HPV-related cancers (HPV+ cancers) continue to contribute to approximately 5% of the worldwide cancer burden [5–14]. HPV 16 is responsible for the majority of HPV+ cancers, contributing to 54% of cervical cancers and ∼90% of HPV+ oropharyngeal squamous cell carcinoma (HPV+OPC) [3,5,8–10,15–17]. While these HPV+ cancers remain prevalent, the majority of total infections are asymptomatic, self-limiting, and clear before cancer progression [3,18–23]. Persistent HPV infection is a necessary component of cancer development but is not considered sufficient without additional co-factors [15,24]. One key factor in maintaining viral persistence is the ability of HPV to evade host immunity [22,23,25–29]. Numerous studies have demonstrated that HPV suppresses innate immune-related signaling in both infected epithelia and neighboring stromal fibroblasts [22,23,25–36]. Suppression of immune-related genes allows for immune evasion, which is critical for viral persistence and may play a role in cancer development [37,38].

The stroma is a complex connective tissue comprised of numerous cell types; the main component of the dermal stroma is fibroblasts [15,39–41]. Fibroblasts support tissue homeostasis via the secretion of all components of the extracellular matrix (ECM) and facilitate stromal extracellular signaling; factors produced by fibroblasts are key for angiogenesis, inflammation, wound healing, and are necessary for the proper differentiation of keratinocytes [23,41,42]. Keratinocyte differentiation is critical for the HPV lifecycle [43,44]. While HPV exclusively infects basal keratinocytes, viral gene products alter the secretion of host factors, indirectly affecting neighboring keratinocytes, fibroblasts and immune cells in the local microenvironment [22,23,45]. Given the complexity of the tissue infected and the transformation process, the relationship between HPV and epithelial-stromal communication remains at a nascent phase and further investigations are warranted [15,23].

The importance of stromal support in the microenvironment is now an emerging field in the context of overall cancer progression, as well as HPV-induced transformation and carcinogenesis [15,23,26,39,40,45–56]. Precise mechanisms for viral transformation and progression mechanisms remain unclear; however, persistent viral oncogene expression contributes to clear epithelial growth advantages [27,57–59,59–64]. HPV E6 and E7 are considered the major viral oncoproteins that contribute to carcinogenesis via altering cellular tumor suppressor pathways; E6 targets and degrades p53, while E7 targets and degrades retinoblastoma protein (pRb) [18,37,57,65,66]. The lesser characterized minor oncoprotein, HPV E5, appears to regulate cellular transformation, immune modulation, and response to cell signaling events [23,57,67]. While the expression of E6 and E7 extends the proliferative capacity of epithelial cells, fibroblasts have demonstrated a cooperative role in the induction of cell immortalization [15,68–71]. E5 has also demonstrated regulatory interactions as an innate immune suppressor in the adjacent stroma, thus contributing to viral persistence [22,23]. Of note, the viral DNA binding protein, E2, is not proposed to be oncogenic but has also been reported to be involved in the suppression of the innate immune response and is crucial for viral episome persistence [28,29,72–75].

Oncogene expression alone is considered insufficient for carcinogenesis, and other indeterminate events have been implicated in transformation [76]. During the HPV lifecycle, the viral genome exists in an episomal form in basal keratinocytes. Conversely, when aberrant HPV genome integration events occur, they have been noted as contributing factors in transformation; viral integration correlates with increased viral oncogene expression, loss of functional E2, cellular growth advantages, enhanced tumor progressiveness, cervical cancer progression, and poor clinical prognostics of HPV+OPC [25,27,59–61,77–85]. It is generally accepted that HPV+ keratinocyte cell lines must be grown in co-culture with fibroblasts to support viral episome maintenance [80,86,87]. HPV+ keratinocytes maintained in the absence of fibroblasts are noted to quickly integrate or lose viral genome expression [87,88]. From these observations, fibroblasts are influential on the HPV episomal status of adjacent keratinocytes, suggesting their role in regulating this transforming factor. The mechanisms of episomal regulatory control via fibroblasts have yet to be elucidated.

We previously reported the value of fibroblast co-culture both in the context of HPV episomal maintenance and as a model for better predicting *in vitro* to *in vivo* translational treatment paradigms [88]. In our previous analysis, we demonstrated that mitomycin C (MMC) growth-arrested murine 3T3-J2 fibroblasts (referred to as J2s moving forward) supported HPV16 long control region (LCR) transcriptional regulation [88]. We further investigated HPV protein expression and host protein signaling observed in the presence or absence of J2s [88]. N/Tert-1 cells (telomerase immortalized foreskin keratinocytes, HPV negative), HFK+E6E7 (foreskin keratinocytes immortalized by the viral oncogenes only), and HFK+HPV16 (foreskin keratinocytes immortalized by the entire HPV16 genome, replicating as an episome), were cultured in the presence or absence of J2s. We demonstrated that HFK+HPV16 maintained in J2 had measurable E7 protein levels; however, when J2s were removed for one week, E7 protein expression was lost [88]. Conversely, there were no significant alterations in E7 protein levels in HFK+E6E7 in the presence or absence of J2s, suggesting a partial reliance on the expression of the LCR or the full genome for the ability of fibroblasts to regulate viral protein expression [88]. Alterations in the protein levels of p53, pRb, and γH2AX were also demonstrated to be altered in the presence of J2 and further suggested fibroblasts may alter host protein expression that is supportive of HPV viral genome regulation [88].

In this report, we utilized RNA sequencing (RNA-seq) and proteomic analysis for a global and comprehensive approach to investigate keratinocyte signaling impacted by fibroblasts. Our investigation confirmed the prior observation, that HPV downregulates portions of innate immune signaling [23,28,29,89–91]. Further separation of keratinocytes grown in the presence or absence of J2s revealed the novel observation that fibroblasts impact the transformation potential of keratinocytes. N/Tert-1+HPV16 cells grown with J2s showed a gene regulation pattern similar to that of a suprabasal layer. Gene ontology (GO) analysis indicated that fibroblasts supported the viral life cycle, and that keratinocytes were less transformed compared to those grown without J2s. In contrast, N/Tert-1+E6E7 cells grown with J2s showed a greater tendency toward transformation than those grown without J2s, especially in relation to altered cell cycle regulation, and oncogenic cytokine expression. Proteomic analysis further supported these observations. Our results confirm that the expression of episomal HPV is necessary to regulate optimal viral-host interactions. Integration would mimic results observed in N/Tert-1+E6E7 cells, and the presence of fibroblasts promote a much more transformed genotype. Overall, our findings suggest that both monoculture and fibroblast co-culture approaches are useful for future studies on HPV-related transformation.

## 2 Materials and methods

### 2.1 Cell Culture

N/Tert-1 cells and all derived cell lines have been described previously and were maintained in keratinocyte-serum free medium (K-SFM; Invitrogen), and supplemented with previously described antibiotics [27–29,88,92–95].

### 2.2 Culture and mitomycin C (MMC) inactivation of 3T3-J2 mouse embryonic fibroblast feeder cells, and co-culture with keratinocytes

As previously described, 3T3-J2 immortalized mouse embryonic fibroblasts (J2) were grown in DMEM and supplemented with 10% FBS [88]. 80-90% confluent plates were supplemented with 4µg/ml of MMC in DMSO (Cell Signaling Technology) for 4-6 hours at 37°C. MMC-supplemented medium was removed and cells were washed with 1xPBS.

Cells were trypsinized, centrifuged at 800 rcf for 5 mins, washed once with 1xPBS, centrifuged again, and resuspended at 2 million cells per mL. Quality control of inactivation (lack of proliferation) was monitored for each new batch of mitomycin-C. Unless otherwise stated, 100-mm plate conditions were continually supplemented with 1x10^6^ J2 every 2-3 days. Before trypsinization or harvesting, plates were washed to remove residual J2.

### 2.3 RNA isolation

The SV total RNA isolation system kit (Promega) was utilized to isolate RNA from cells, as per the manufacturer’s protocol.

### 2.4 Human Sequences RNA-seq Bioinformatics Pipeline

Library preparation, sequencing, and pre-processing of samples was performed by Novogene. Novogene uses in-house scripts to clean raw reads, filtering out low-quality reads, and reads containing adapter sequences. The genome index was built and cleaned sequences were aligned to the reference human genome using Hisat2 v2.05 [96,97]. Raw gene expression levels were quantified with featureCounts v1.5.0-p3 and then normalized to fragments per kilobase per million (FPKM) [98]. Differential expression analysis was performed using DESeq2 R package v1.20.0 between three experimental groups N/Tert-1, N/Tert-1+E6/E7, and N/Tert-1+HPV16 treated with J2 fibroblasts (n=3 in each group) and their paired controls respectively (untreated). P-values were adjusted using the Benjamini and Hochberg’s approach for controlling the false discovery rate (FDR), where significance for a differentially expressed gene was determined at FDR < 0.05 [99].

### 2.4 Gene Ontology Enrichment Analysis

GO enrichment analysis of differentially expressed genes was implemented by the clusterProfiler R package, in which gene length bias was corrected [100,101]. GO terms with corrected P-value < 0.05 were considered significantly enriched by differential expressed genes. Heatmaps were generated with the ’pheatmap’ R package using z-score normalized FPKM gene expression averages for each sample condition.

### 2.5 HPV16 sequences RNA-seq Bioinformatics Pipeline

Fastq files from Novogene were examined for quality using FastQC and quality control reports were collated by multiQC [102,103]. Reads were filtered to remove low quality sequences and adapter sequences were trimmed using trimmomatic v 0.39 [104]. A genome index was built and all sequences were aligned to the GRCh38.d1.vd1 Reference Sequence, part of the Genomic Data Commons GDC data harmonization pipeline, using STAR aligner v 2.7.9.a [105]. Samtools v1.16.1 was used to index and filter the bam file for reads aligned to HPV16 [106]. The HPV16 filtered bam files were converted back to fastq files using bedtools [107]. The HPV16 fastq sequences were re-aligned to an HPV16 reference genome from NCBI and raw gene expression levels were counted using featureCounts. Raw counts were then normalized using EdgeR’s calcNormFactors scaling factor of trimmed mean of M-values (TMM) normalization. EdgeR’s quasi-likelihood F-test (QLF) method was then used for differential expression analysis of each gene between three experimental groups N/Tert-1, N/Tert-1+E6/E7, and N/Tert-1+HPV16 treated with J2 fibroblasts (n=3 in each group) and their paired controls respectively (untreated) [108–110]. The p-value of each QLF test was adjusted using a Benjamini-Hochberg False Discovery Rate (FDR) multiple testing correction using the basic R stats package p.adjust function. Genes passing the FDR cut-off threshold of ≤ 0.05 for significance were considered statistically significantly different.

### 2.6 Real-time PCR (qPCR)

A high-capacity cDNA reverse-transcription kit from Invitrogen was used to synthesize cDNA from RNA and processed for qPCR. qPCR was performed on 10 ng of the cDNA isolated. cDNA and relevant primers were mixed with PowerUp SYBR green master mix (Applied Biosystems), and real-time PCR was performed using the 7500 Fast real-time PCR system, using SYBR green reagent. Expression was quantified as relative quantity over GAPDH using the 2−ΔΔCT method. Primer used are as follows. HPV16 E2 F, 5′-ATGGAGACTCTTTGCCAACG-3′; HPV16 E2 R, 5′-TCATATAGACATAAATCCAG-3′; HPV16 E6 F, 5′-TTGAACCGAAACCGGTTAGT- 3′; HPV16 E6 R, 5′-GCATAAATCCCGAAAAGCAA-3′; MX1 F, 5′- GGTGGTCCCCAGTAATGTGG-3′; MX1 R, 5′-CGTCAAGATTCCGATGGTCCT-3′; STAT1 F, 5′-CAGCTTGACTCAAAATTCCTGGA-3′; STAT1 R, 5′- TGAAGATTACGCTTGCTTTTCCT-3′; STAT2 F, 5′-CCAGCTTTACTCGCACAGC- 3′; STAT2 R, 5′-AGCCTTGGAATCATCACTCCC-3′; STAT3 F, 5′- CAGCAGCTTGACACACGGTA-3′; STAT3 R, 5′- AAACACCAAAGTGGCATGTGA-3′; p53 F, 5′-GAGGTTGGCTCTGACTGTACC-3′; p53 R, 5′-TCCGTCCCAGTAGATTACCAC-3′; Glyceraldehyde-3-phosphate dehydrogenase (GAPDH) F, 5′-GGAGCGAGATCCCTCCAAAAT-3′; GAPDH R, 5′- GGCTGTTGTCATACTTCTCATGG-3′.

### 2.7 Exo V

PCR based analysis of viral genome status was performed using methods described by Myers *et al.* [111]. 20 ng of genomic DNA was either treated with exonuclease V (RecBCD, NEB), in a total volume of 30 ul, or left untreated for 1 hour at 37°C followed by heat inactivation at 95°C for 10 minutes. 2 ng of digested/undigested DNA was then quantified by real time PCR, as noted above, using and 100 nM of primer in a 20 μl reaction. Nuclease free water was used in place of the template for a negative control. The following cycling conditions were used: 50°C for 2 minutes, 95°C for 10 minutes, 40 cycles at 95°C for 15 seconds, and a dissociation stage of 95°C for 15 seconds, 60°C for 1 minute, 95°C for 15 seconds, and 60°C for 15 seconds. Separate PCR reactions were performed to amplify HPV16 E6 F: 5’- TTGCTTTTCGGGATTTATGC-3’ R: 5’- CAGGACACAGTGGCTTTTGA-3’, HPV16 E2 F: 5’- TGGAAGTGCAGTTTGATGGA-3’ R: 5’- CCGCATGAACTTCCCATACT-3’, human mitochondrial DNA F: 5’-CAGGAGTAGGAGAGAGGGAGGTAAG-3’ R: 5’- TACCCATCATAATCGGAGGCTTTGG -3’, and human GAPDH DNA F: 5’- GGAGCGAGATCCCTCCAAAAT-3’ R: 5’- GGCTGTTGTCATACTTCTCATGG-3’

### 2.8 Proteomic sample preparation

The samples were digested using commercially available PreOmics iST sample clean up protocol. To the sample containing approximately 100ug of protein, 70ul of lysis buffer was added and mixed, followed by an incubation for 10 minutes at 950C; 1000rpm. 50ul of DIGEST solution was added to the mixture, which was then incubated at 370C for 3hrs at 500 rpm. After the digestion, 100ul of STOP solution was added and mixed properly. The digest was then centrifuged at 3800rcf; 3min to ensure complete flow through and washed with 200ul of WASH 1 and 200ul of WASH 2 solution followed by centrifugation after each wash. The cartridge was then placed to the fresh collection tube and 100ul of ELUTE solution was added and centrifuged at 3800rcf; 3min to ensure complete flow through. This step was repeated one more time to ensure maximum recovery. The elutes were then placed in a vacuum evaporator at 450C until completely dried.

### 2.9 LC-MS/MS

LC-MS/MS analysis were performed using a Q-Exactive HF-X (Thermo) tandem mass spectrometer coupled to an Easy nLC 1200 (Thermo) nanoflow UPLC system. The LC-MS/MS system was fitted with an Easy spray ion source and an Acclaim PepMap 75µm x 2cm nanoviper C18 3µm x 100Å pre-column in series with an Acclaim PepMap RSLC 75µm x 50cm C18 2µm bead size (Thermo). The mobile phase consists of Buffer A (0.1% formic acid in water) and Buffer B (80% acetonitrile in water,0.1% formic acid). 500ng of peptides were injected onto the above column assembly and eluted with an acetonitrile/0.1% formic acid gradient at a flow rate of 300 nL/min over 2 hours. The nano-spray ion source was operated at 1.9 kV. The digests were analyzed using a data dependent acquisition (DDA) method acquiring a full scan mass spectrum (MS) followed by 40 tandem mass spectra (MS/MS) in the high energy C-trap Dissociation HCD spectra). This mode of analysis produces approximately 50,000 MS/MS spectra of ions ranging in abundance over several orders of magnitude. Not all MS/MS spectra are derived from peptides.

### 2.10 Proteomic Data Analysis

The data were analyzed in Proteome Discoverer (ver 3.0) using the Sequest HT search algorithm and the Human database. Proteins were identified at an FDR < 0.01 and quantification used the peptide intensities. Raw protein abundances were normalized in Proteome Discoverer using the “Total Peptide Abundance” method. Differential Enrichment of protein abundance was performed using the ’DEP’ package v. 1.26 [112]. First, we filtered for proteins detected in two of three replicates of at least one of the experimental conditions. Variance stabilizing transformation of remaining protein intensity observations was performed using the ’vsn’ package v 3.72 via the ’normalize_vsn’ function [113]. The quantile regression-based left-censored (QRILC) method was used as the missing value imputation approach. The differential enrichment test was conducted pairwise on each protein using limma v 3.60.4 between three experimental groups N/Tert-1, N/Tert-1+E6/E7, and N/Tert-1+HPV16 treated with J2 fibroblasts (n=3 in each group) and their paired controls (untreated), respectively [114]. Proteins were identified as significantly differentially expressed between the control and experimental groups with a Benjamini-Hochberg adjusted p-value of < 0.05, and a |log2-fold change| > 0.58.

### 2.11 Immunoblotting

Cells were trypsinized, washed with PBS and resuspended in 2x pellet volume NP40 protein lysis buffer (0.5% Nonidet P-40, 50 mM Tris [pH 7.8], 150 mM NaCl) supplemented with protease inhibitor (Roche Molecular Biochemicals) and phosphatase inhibitor cocktail (MilliporeSigma). Cell suspension was incubated on ice for 20 min and then centrifuged for 20 min at 184,000 rcf at 4 °C. Protein concentration was determined using the Bio-Rad protein estimation assay according to manufacturer’s instructions. 50 μg protein was mixed with 2x Laemmli sample buffer (Bio-Rad) and heated at 95 °C for 5 min. Protein samples were separated on Novex 4–12% Tris-glycine gel (Invitrogen) and transferred onto a nitrocellulose membrane (Bio-Rad) at 30V overnight using the wet-blot transfer method. Membranes were then blocked with Odyssey (PBS) blocking buffer (diluted 1:1 with PBS) at room temperature for 1 hr. and probed with indicated primary antibody diluted in Odyssey blocking buffer, overnight. Membranes were washed with PBS supplemented with 0.1% Tween (PBS-Tween) and probed with the Odyssey secondary antibody (goat anti-mouse IRdye 800CW or goat anti-rabbit IRdye 680CW) (Licor) diluted in Odyssey blocking buffer at 1:10,000. Membranes were washed twice with PBS-Tween and an additional wash with 1X PBS. After the washes, the membrane was imaged using the Odyssey^®^ CLx Imaging System and ImageJ was used for quantification, utilizing GAPDH as internal loading control. Primary antibodies used for western blotting studies are as follows: pRb 1:1000 (Santa Cruz, sc-102), p53 1:1000 (Cell Signaling Technology, CST-2527, and CST-1C12), γH2AX 1:500 (Cell Signaling Technology, CST-80312 and CST-20E3).

### 2.12 Reproducibility, research integrity, and statistical analysis

All experiments were carried out at least in triplicate in all of the cell lines indicated. Keratinocytes were typed via cell line authentication services. All images shown are representatives from triplicate experiments. Student’s t-test or analysis of variance was used to determine significance as appropriate: *P < 0.05, **P < 0.01, ***P < 0.001.

## 3 Results

### 3.1 Differential Genomic Landscapes altered by fibroblasts in keratinocytes

The utility of a supportive fibroblast feeder layer is broadly accepted as essential for maintaining an episomal HPV genome in primary keratinocyte models, and is a necessary component of 3D models for HPV lifecycle analysis where it is chiefly responsible for proper keratinocyte differentiation [57,68,77,87,88,115–122]. While the coculture of keratinocytes with fibroblast feeders is accepted, the full mechanism of how fibroblasts aid in HPV episomal maintenance has yet to be deciphered. It is worth noting that 2D coculture may represent interactions that occur in the basal layer, while far more complex spatial and temporal regulatory mechanisms are likely involved in 3D models and *in vivo*. This analysis focuses on short-term 2D interactions, with the aim of investigating 3D models in the future.

We previously demonstrated that fibroblast co-culture was important for maintaining HPV episomes, influenced HPV16 LCR transcriptional regulation, and supported the expression of HPV16 E7 protein in human foreskin keratinocytes immortalized with HPV16 (HFK+HPV16) [88]. We also observed that fibroblasts altered host protein levels which could affect viral genome regulation [88]. Taking a more global approach to investigate signaling impacted by fibroblasts, N/Tert-1, N/Tert-1+E6/E7, and N/Tert-1+HPV16 cells were cultured in the presence or absence of J2s for one week. These matched samples were then subjected to bulk RNA-seq analysis, and label-free liquid chromatography-mass spectrometry-based proteomic analysis (LC-MS/MS).

For RNA-seq, triplicate sample data were combined to assess differential gene expression analysis. Initial comparisons were made in large batched sets; cell lines were either not separated based on the presence or absence of J2, or grouped as all mono-culture vs all co-culture. They were compared in the following large sets: N/Tert-1 vs N/Tert-1+HPV16, N/Tert-1+E6E7 vs N/Tert-1+HPV16, N/Tert-1 vs N/Tert-1+E6E7, and monoculture control vs co-culture “+J2”. Evaluations of datasets were then further compared based on the presence or absence of J2 in each individual N/Tert-1, N/Tert-1+E6E7, or N/Tert-1+HPV16 cell line and cross-compared. Our data revealed numerous genes significantly differentially expressed 1.5 fold or greater when cross-comparing our samples (DEG gene counts presented in Figure 1A, Quantitative correlation presented in Figure 1B). A full list of these genes can be found in Supplementary Material S1. The expression level of the HPV16 genes used to generate the gene expression data is given in Supplementary Table S2. Novogene and further bioinformatic analysis identified the most affected canonical pathways, upstream regulators, diseases, and functions predicted to be altered in this data set; significant observations are given in Supplementary Tables S3. The most notable HPV differential expression and GO enrichment observations were alterations in innate immune signaling, including altered cytokine and chemokine activity; additional alterations in cellular communication potential, tight junction regulation, and growth factor signaling events were also differentially regulated (GO enrichment plots summarized in Figures 2A-C). When grouped as a whole, fibroblasts significantly altered GO enrichment associated with angiogenesis, differentiation, extracellular matrix organization, and both cytokine and growth factor-related activity (Figure 2D).

**Figure 1.**
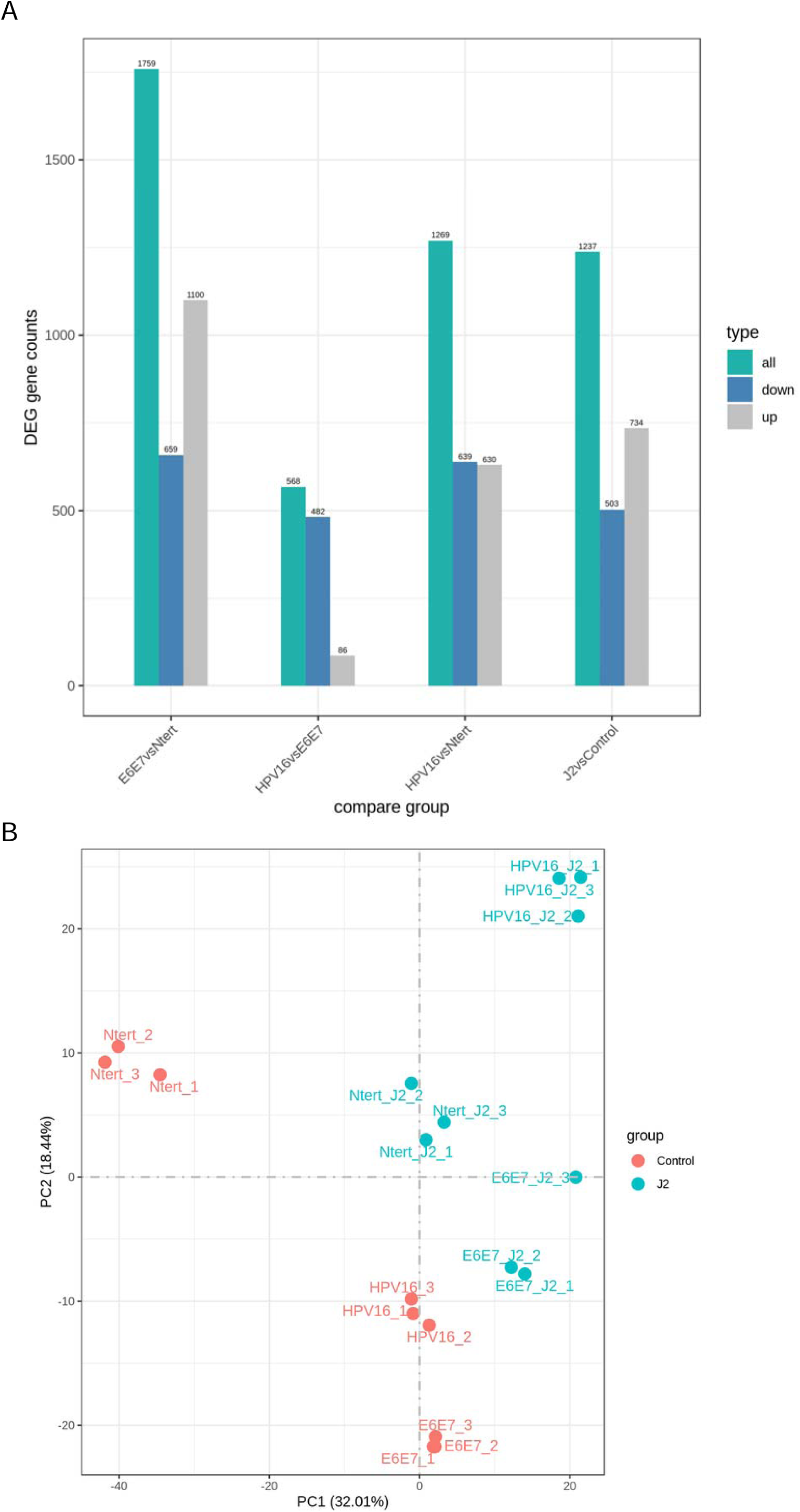
Global comparison of RNA-seq. **1A.** RNA-seq differential expression (DEG) analysis histogram comparison of the number of significant differential genes (including up-regulation and down-regulation) for each combination. **1B.** Principal component analysis (PCA) analysis on the gene expression value (FPKM) of all samples.

**Figure 2.**
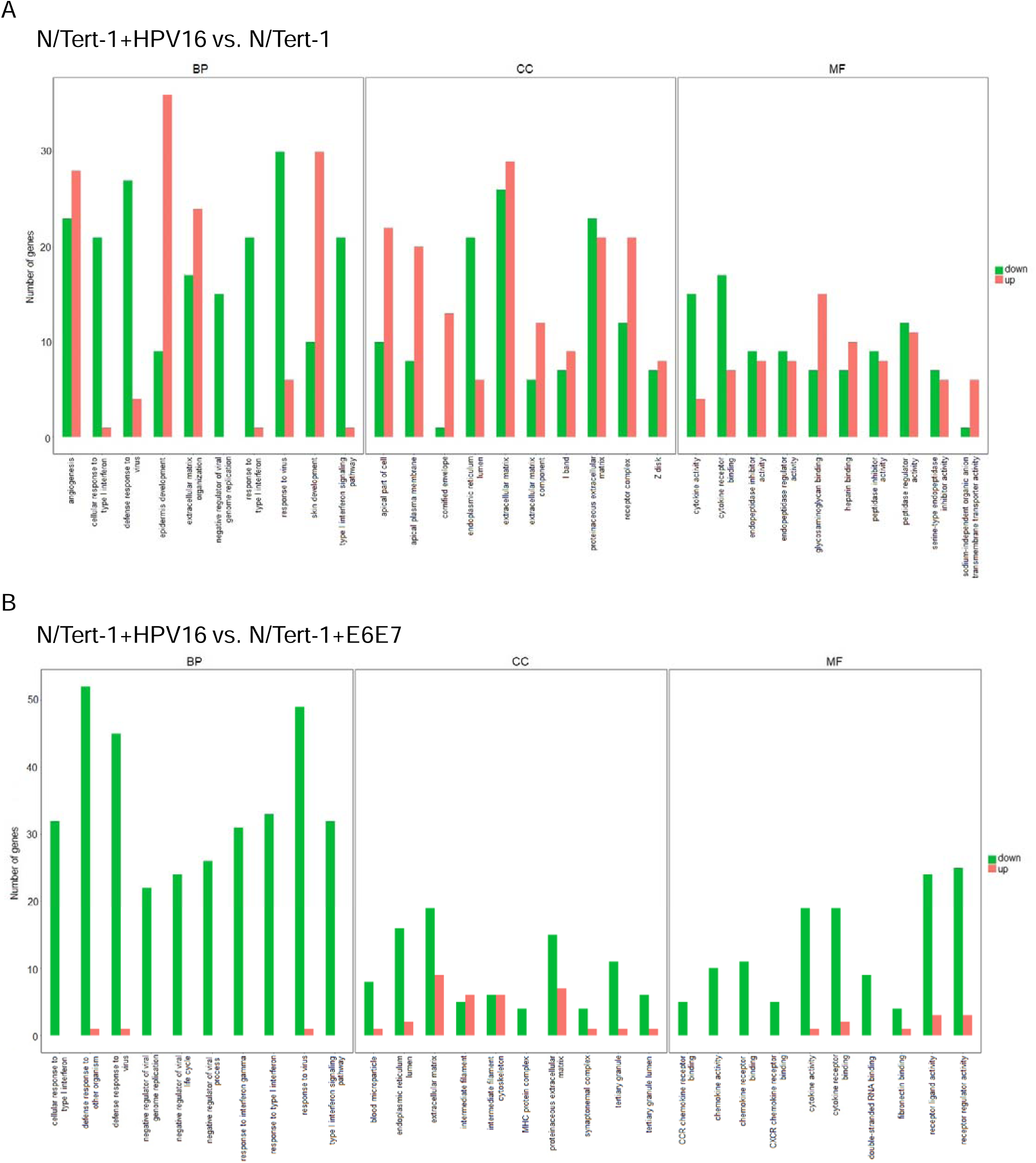
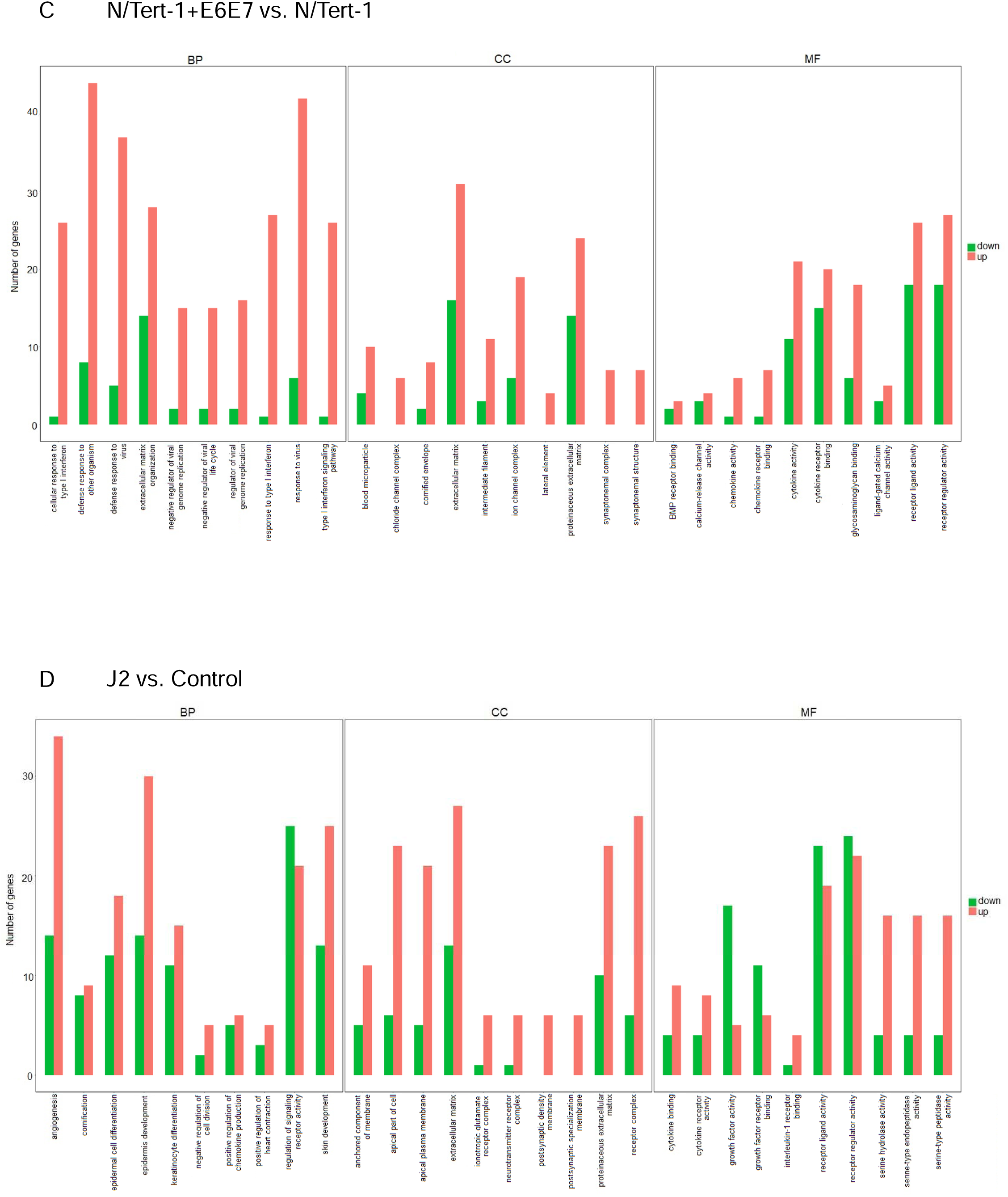
Gene ontology (GO) enrichment analysis histograms demonstrate differential regulation between N/Tert-1 cell lines and between mono vs co-culture. The 30 most significantly GO terms are displayed. All Terms are separated according to major categories of biological processes (BP), cell components (CC), molecular functions (MF) and categories of upregulated and downregulated expression of noted GO. **2A.** Grouped N/Tert-1+HPV16 are compared to Grouped N/Tert-1. **2B.** Grouped N/Tert-1+HPV16 are compared to Grouped N/Tert-1+E6E7. **2C.** Grouped N/Tert-1+E6E7 are compared to Grouped N/Tert-1. **2D.** Grouped fibroblast co-culture cell line sets (J2) are compared to Grouped mono-culture cell line sets (Control).

As previously reported, numerous gene sets related to interferon (IFN) response were significantly reduced in the N/Tert-1+HPV16 group, over that of both N/Tert-1 and N/Tert-1+E6E7 groups (Figures 2A-B) [28,123]. Of note, fibroblasts were not utilized when preparing our N/Tert-1-related cultures in previous RNAseq analysis [28,29]. Various interleukins and CXCL family members were also significantly downregulated in grouped N/Tert-1+HPV16 when compared to grouped N/Tert-1 and N/Tert-1+E6E7 (Figures 2A-B). Reactome enrichment further highlighted the following genes concerning the aforementioned significantly downregulated networks: *BST2, CREB5, CSF1, CX3CL1, CXCL1, CXCL2, CXCL3, IFI27, IFI35, IFI6, IFIT1, IFITM1, IFITM3, IL18R1, IL6, IRF7, ISG15, HLA-B, LIF, MMP9, MX1, MX2, OAS1, OAS2, OAS3, PIK3R3, PTAFR, RIPK3, RSAD2, SAMHD1, STAT1, TRIM22, UBE2L6, USP18, XAF1* (Supplemental Tables S3). The observation that HPV downregulates innate immune functions is not novel, but highlights the consistency of our observations with others [23,28,29,89–91].

Several interesting significant alterations in GO enrichment were observed when N/Tert-1 cell lines were further separated based on the presence or absence of J2. N/Tert-1+HPV16 continuously maintained in J2 co-culture demonstrated significant upregulation of interleukin antagonist genes and genes related to inflammation and cell motility, while expression of IFN-induced genes remained downregulated (Figures 3A-J). Genes related to B-cell recruitment and the compliment pathway, also were enriched in N/Tert-1+HPV16 maintained in J2 (Figures 3A,C). The GO enrichment of N/Tert-1+E6E7 in the presence or absence of J2, in comparison to N/Tert-1+HPV16 in the presence or absence of J2, was markedly different. N/Tert-1+E6E7 grown in the presence of J2 exhibited the most significant increase in GO enrichment of genes related to IFN, indicating that the expression of the full viral genome is necessary for their repression (Figures 3A-J). This would correspond to observations that both E2 and E5 have been tied to the regulation of innate immunity [23,28,29]. While IFN is known to regulate viral infections, IFN-mediated activation of the Janus kinase (JAK)-signal transducer activator of transcription (STAT) has also been associated with cancer progression, including HPV+ cervical cancer [124,125]. Specifically, HPV oncoproteins have previously been shown to activate JAK/STAT [125]. GO enrichment, and qPCR validation demonstrate that N/Tert-1+E6E7 cells cocultured with fibroblasts, markedly upregulate *STAT1,2,3* expression; in comparison, N/tert-1+HPV16 keratinocytes cocultured with fibroblasts have significantly lower expression of these genes (Figures 3E-H).

**Figure 3.**
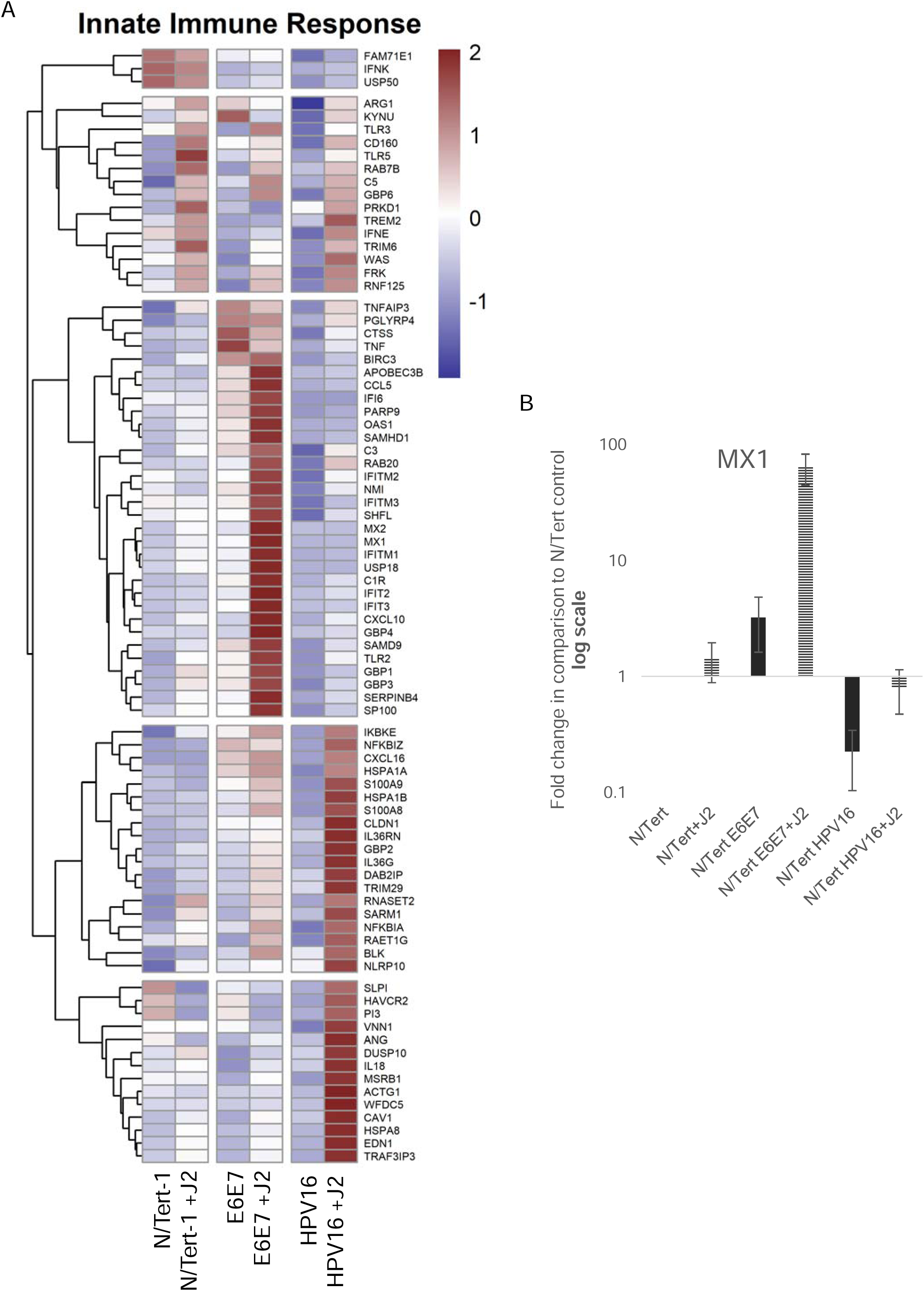
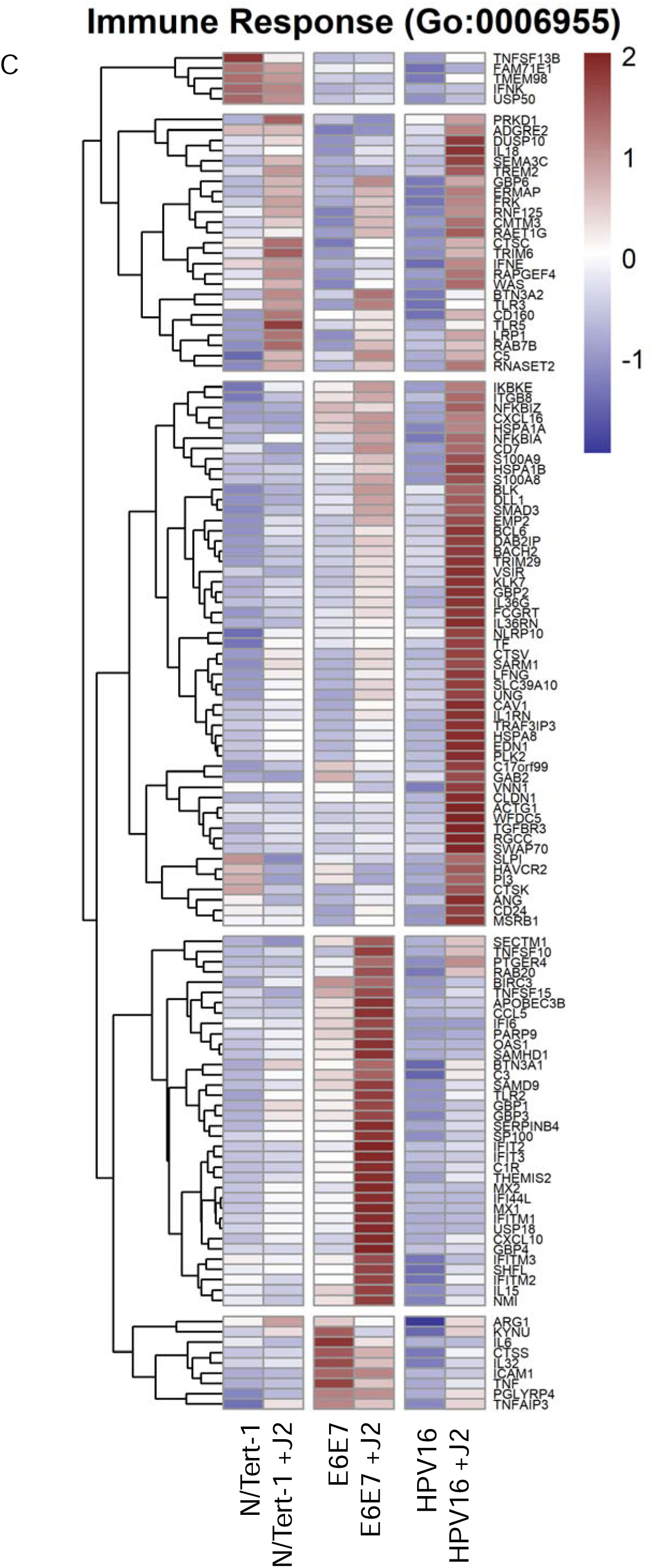
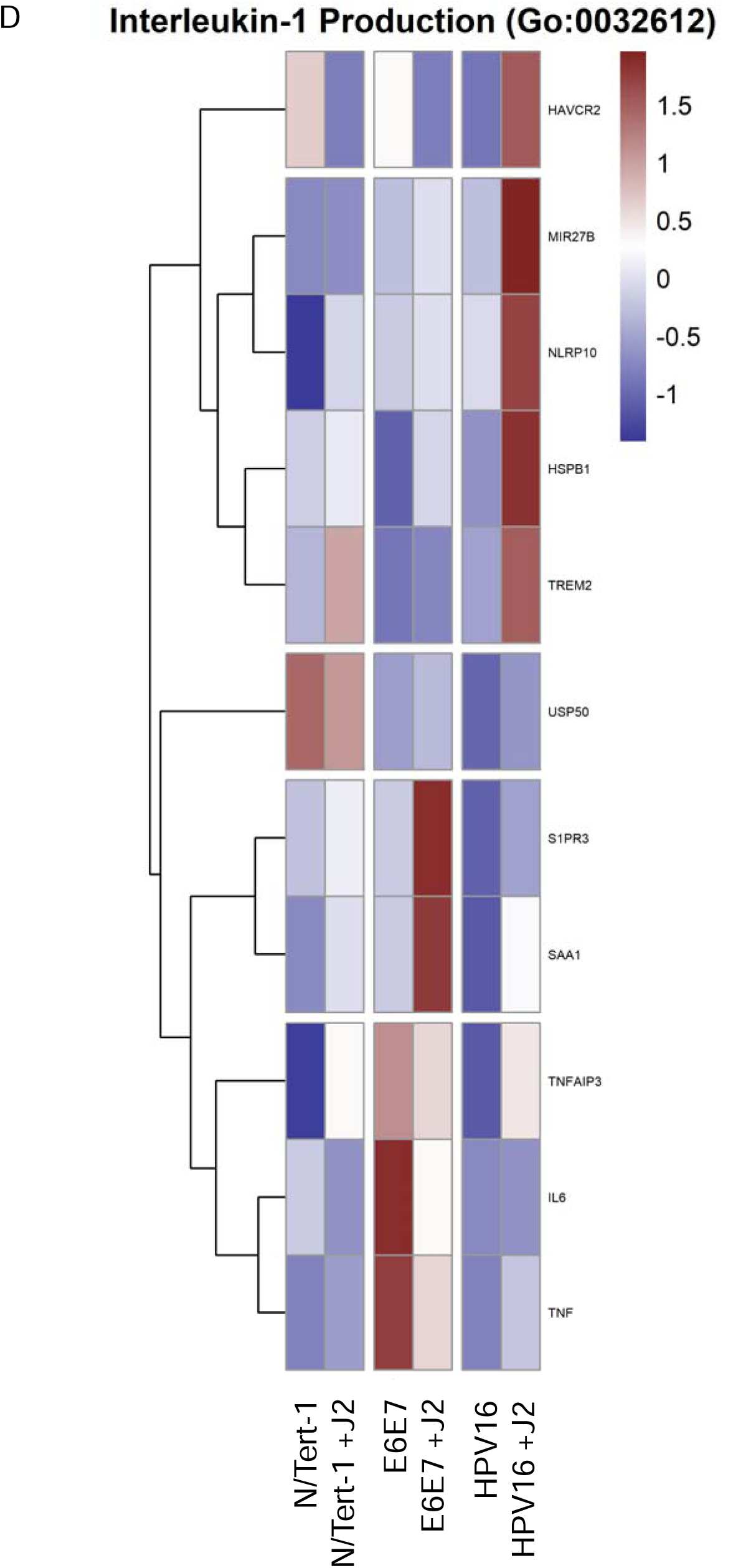
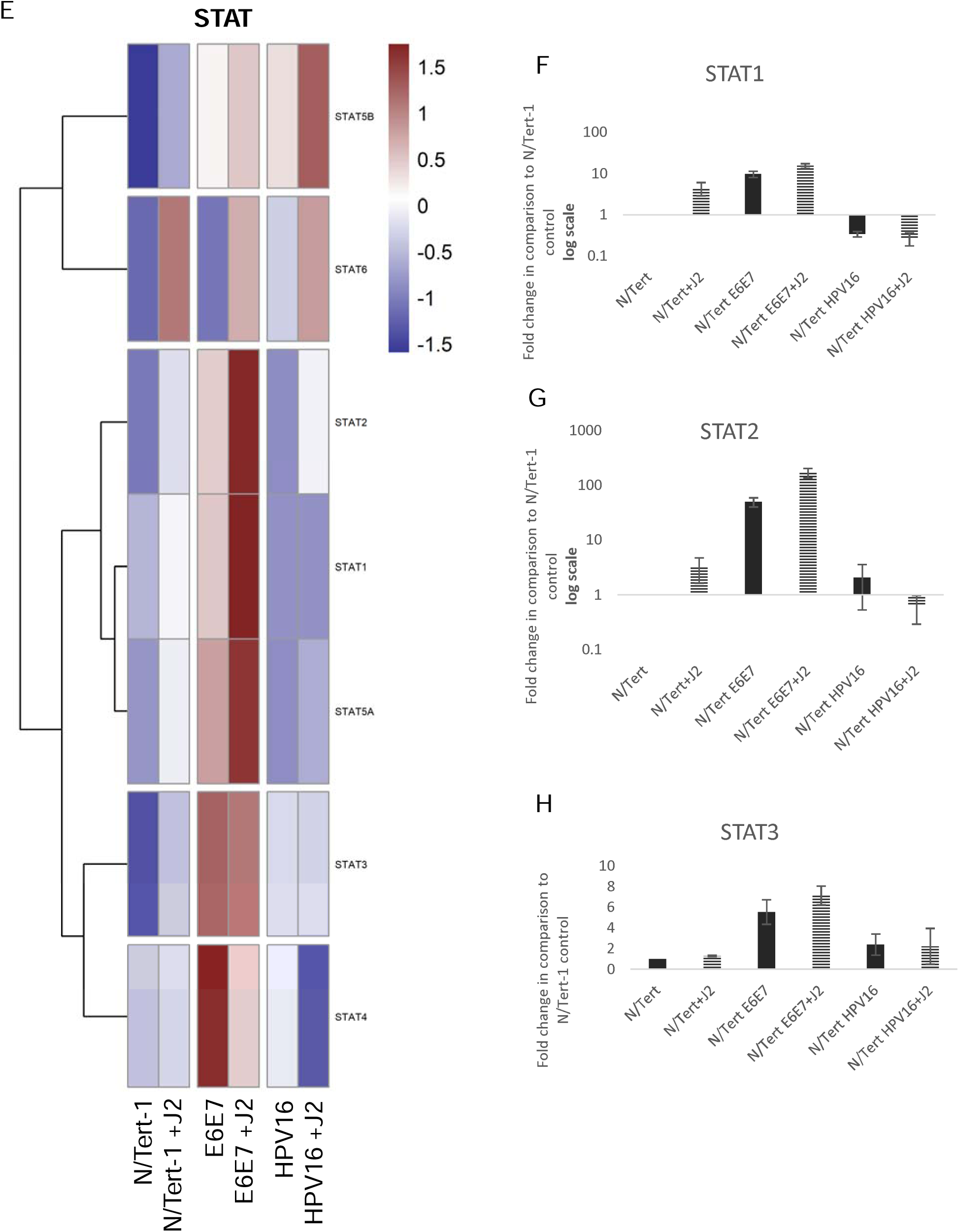
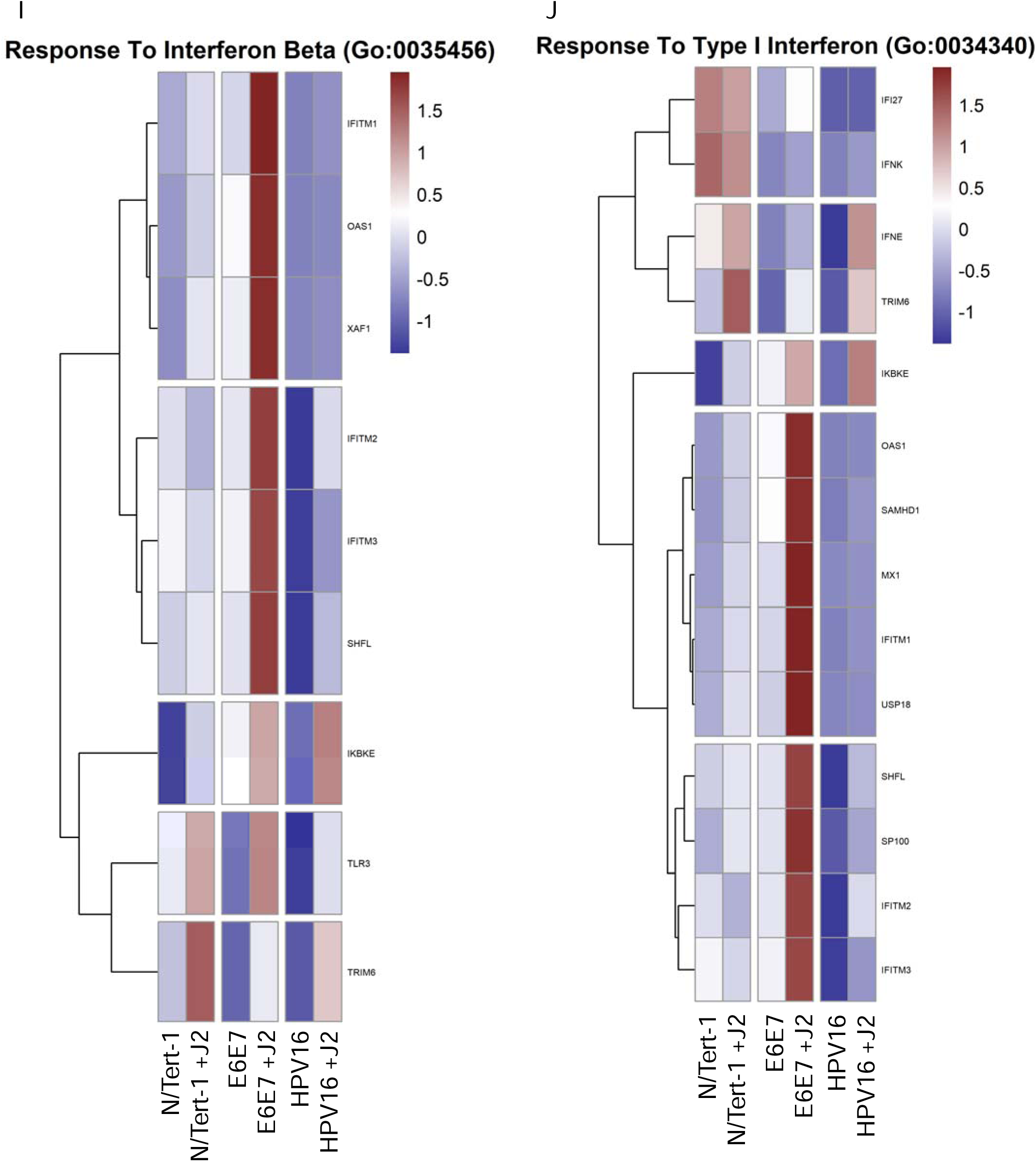
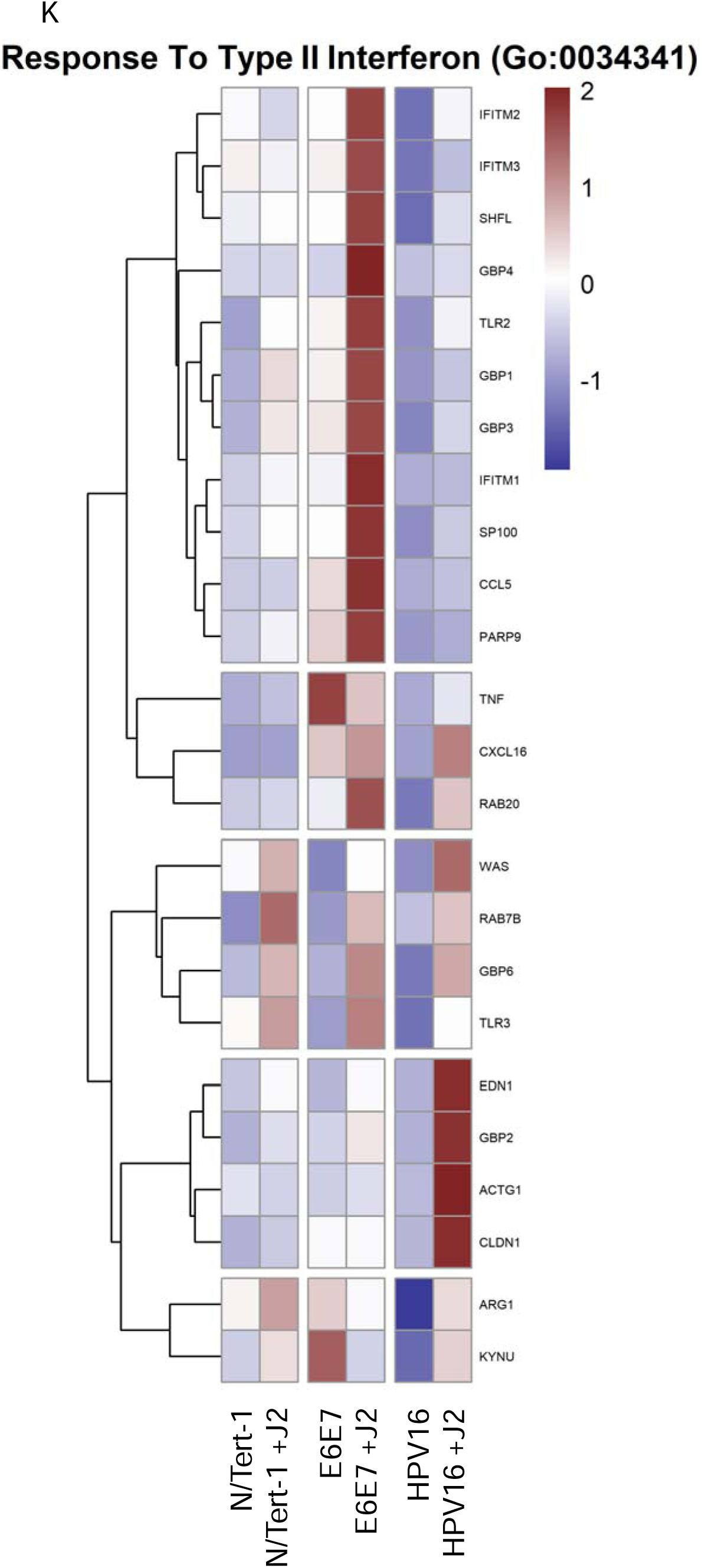
Fibroblasts differentially regulate GO enrichment in relation to innate immune function. **3A.** Heat map demonstrating significant GO:0045087 innate immune regulation across all groups. **3B.** qPCR validation of MX1 RNA expression, presented in log scale. **3C.** Heat map demonstrating significant GO:0006955 innate immune response across all groups. **3D.** Heat map demonstrating significant GO:0032612 interleukin-1 production across all groups. **3E.** Heatmap demonstrating significant STAT RNA expression across all groups. **3F.** qPCR validation of STAT1 RNA expression, presented in log scale. **3G.** qPCR validation of STAT2 RNA expression, presented in log scale. **3H.** qPCR validation of STAT3 RNA expression. **3I.** Heat map demonstrating significant GO:0035456 response to interferon beta across all groups. **3J.** Heat map demonstrating significant GO:0034340 response to type I interferon across all groups. **3K.** Heat map demonstrating significant GO:0034341 response to type II interferon across all groups.

Another noteworthy observation in our GO enrichment cross-comparison, was the alterations observed in genes related to cell junctions, particularly with tight junctions (TJs) and cell-cell signaling control (Figure 4). TJs are comprised of a complex group of molecules, and are associated with the suprabasal and intermediate layers of epithelia. While numerous TJ proteins are downregulated in the transformation process, others are overexpressed and mislocalized [126,127]. Such dysregulation of TJ proteins is associated with epithelial-to-mesenchymal transition (EMT) and invasive phenotypes, including in HPV+ cervical cancer and HPV16 E7 has been shown to alter the expression and localization of TJ-associated claudins [127–129]. Twist1 is also associated with EMT; its transcriptional activation of Claudin-4 has been shown to promote cervical cancer migration and invasion [130–132]. Our analysis shows partial upregulation of TJ components in E6E7+ cells by coculture with fibroblasts, and a significant upregulation in HPV16+ keratinocytes (Figures 4A,C). In particular, there was a marked increase in TJ assembly proteins in both cell lines, including claudins, which are crucial to tight junction integrity (Figure 4A,C). Here, we suggest that this is a model for stages of transformation. The decreased expression of junctional proteins seen in N/Tert-1+E6E7 is more analogous to later, neoplastic stages of transformation; when the viral genome is integrated, E6E7 is overexpressed and there is a progression towards EMT. Meanwhile, the increased expression of TJ components in HPV16+ keratinocytes cultured with fibroblasts is analogous to early viral lifecycle stages. Furthermore, by inducing increased levels of TJ components in infected keratinocytes, the virus induces an environment that mimics a suprabasal phenotype, which is important for the amplification stage of the viral lifecycle [82,118,133]. As large complexes, TJs facilitate signal transduction and are involved in cell proliferation, migration, differentiation, and survival, all of which are beneficial to the viral lifecycle [134]. The comparison to E6E7+ keratinocytes indicates that the upregulation of junctional proteins seen in HPV16+ cells is likely driven by other viral factors, possibly E2, although this warrants further investigation. It would be interesting to further dissect the impact of keratinocyte-fibroblast co-culture upon the subcellular localization of these TJ components and any resulting downstream effects on cell invasive capacity in both E6E7+ and full-genome containing cell lines.

**Figure 4.**
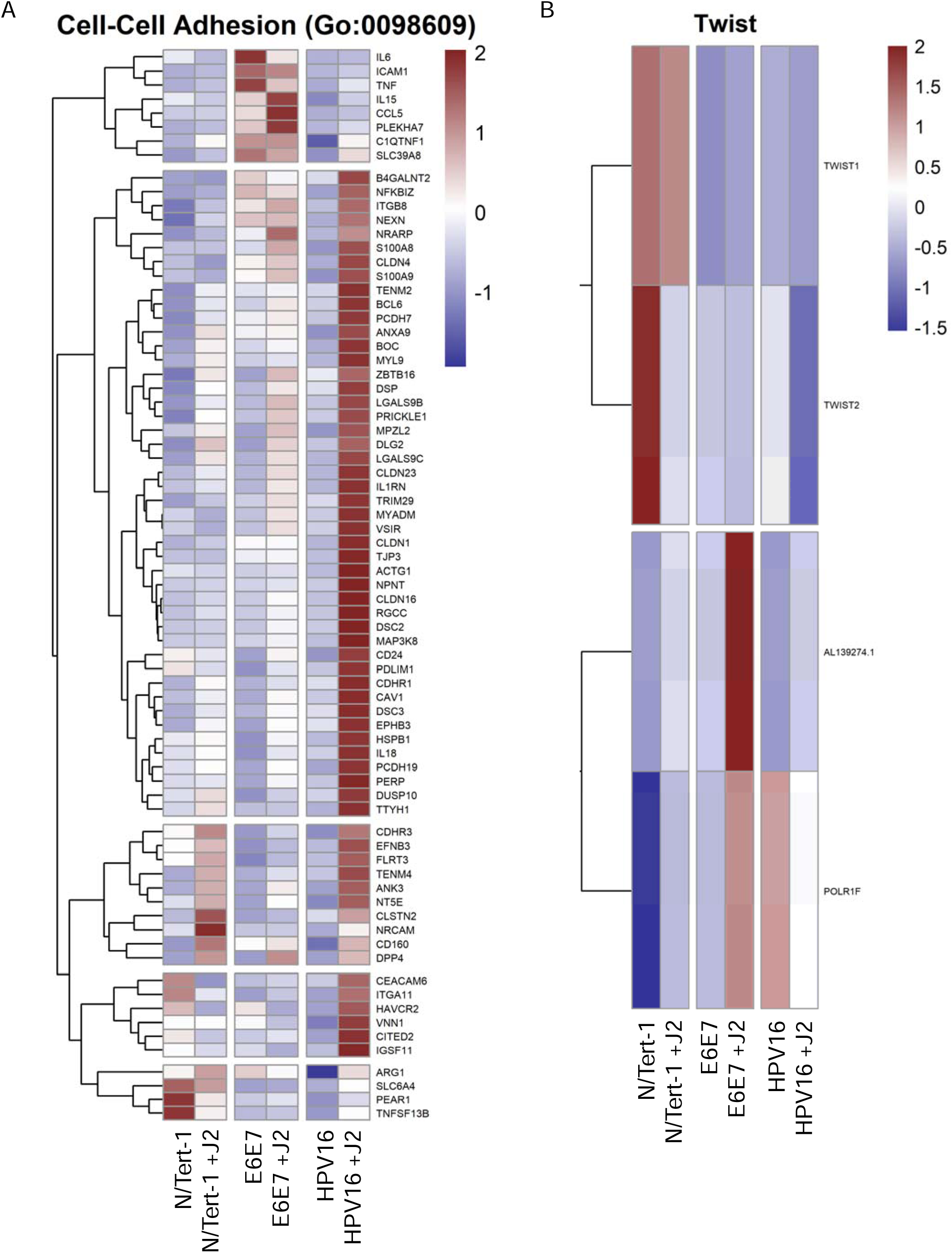
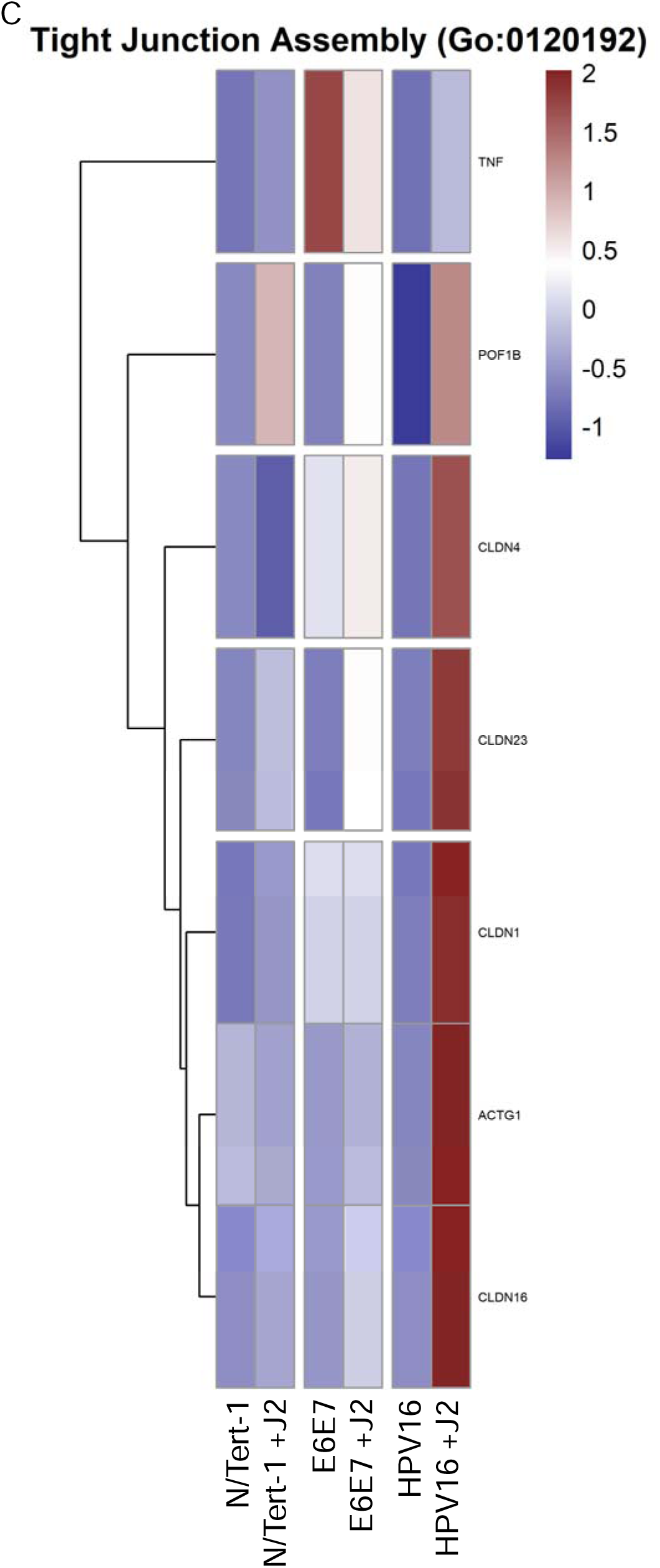
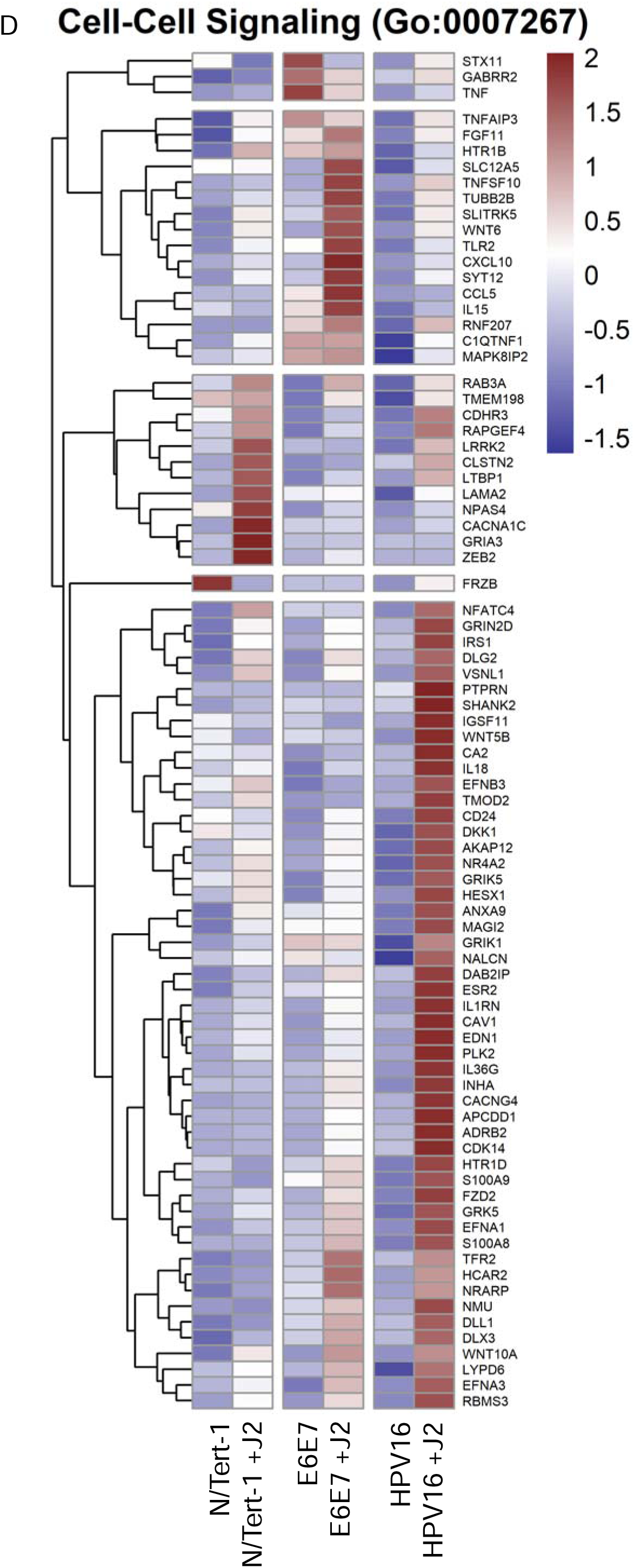
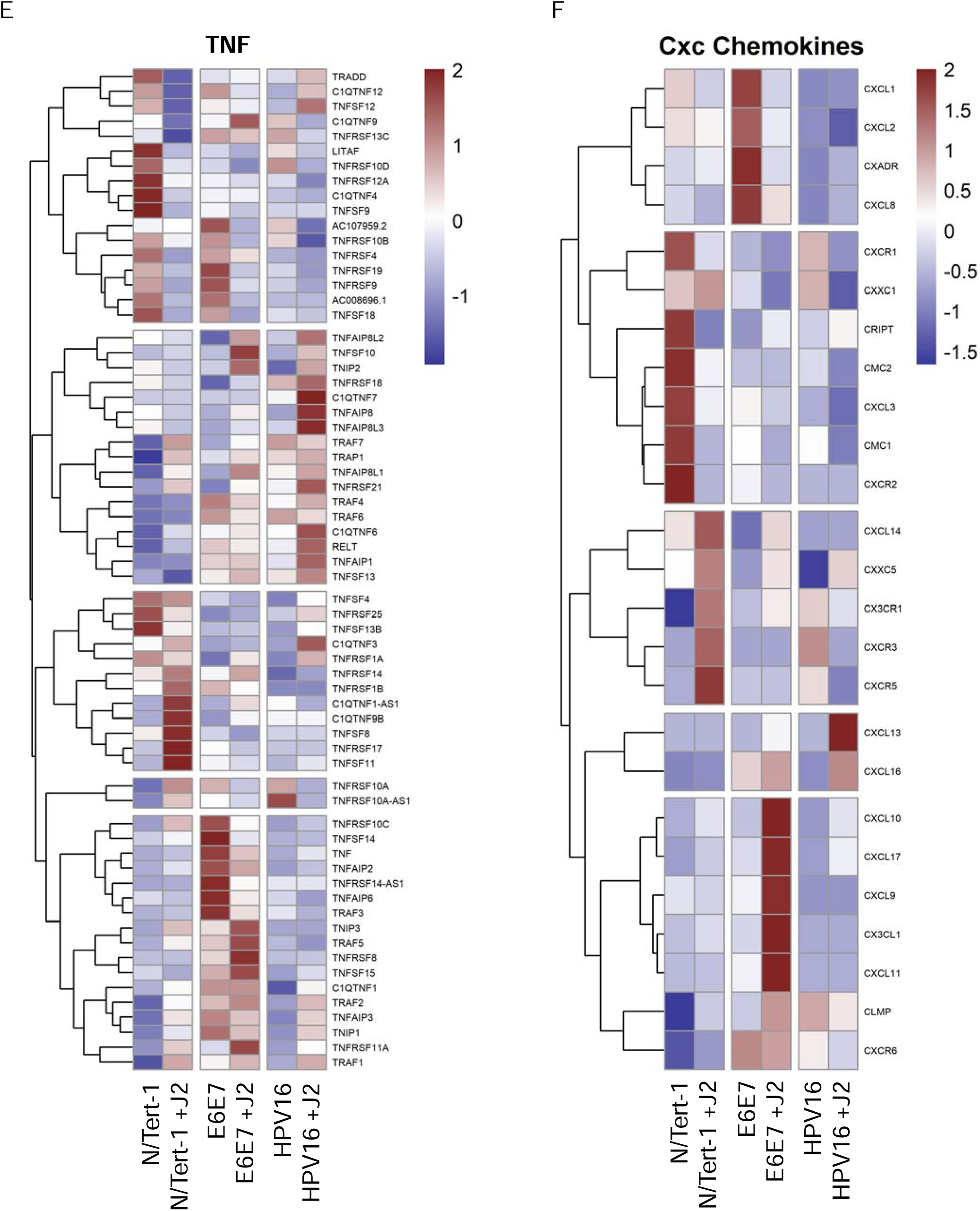
Fibroblasts differentially regulate GO enrichment in relation to cell signaling and epithelial-to-mesenchymal (EMT) progression. **4A.** Heat map demonstrating significant GO:0098609 cell-cell adhesion across all groups. **4B.** Heatmap demonstrating significant TWIST RNA expression across all groups. **4C.** Heat map demonstrating significant GO:0120192 tight junction assembly across all groups. **4D.** Heat map demonstrating significant GO:0007267 cell-cell signaling across all groups. **4E.** Heat map demonstrating significant GO:0033209 TNF across all groups. **4F.** Heat map demonstrating significant CXC chemokines across all groups.

Chemokines are small molecules and secretory peptides are associated with cellular signaling and are broadly divided into subfamilies based on their amino acid motifs: XC, CC, CXC, and CXXXC [135,136]. Chemokine ligands, work jointly with specific chemokine receptors, to control a broad range of biological processes [135,136]. CXC family members are further divided into ELR+ and ELR-members, based on the presence or absence of a Glu-Leu-Arg (ELR) motif in their N-terminus [135]. ELR+ CXC chemokines are associated with the progression of cancer, conversely downregulation of these has been found to suppress the motility of cancer [135]. On the other hand, ELR-CXC chemokines are associated with tumor-suppressive effects [135]. Chemokine-related GO enrichment observed in N/Tert-1+HPV16 grown in the presence of J2 was highly indicative of a less tumorigenic genotype (Figure 4F). This suggests that fibroblasts are likely playing a role in preventing the transformation of HPV+ keratinocytes. Moreover, GO enrichment of *TWIST* expression (Figure 4B) demonstrated that N/Tert-1+HPV16 grown in the presence of J2 is indicative of a less transformed genotype [132,137,138]. CXC-related signaling is known to impact EMT and cancer progression via interactions with β-catenin, TNF, and Notch/Wnt signaling [135,136,139–142]. While these signaling pathways can have both tumor-promoting and suppressive roles that are cancer-dependent, it is clear that fibroblasts are altering the GO enrichment of N/Tert-1+HPV16 grown in the presence of J2, and this has implications in the mechanism of HPV16-driven carcinogenesis (Figure 4).

As we previously observed protein alterations in p53, pRb, and γH2AX in our human foreskin (HFK) cell lines, we also confirmed this trend via western blotting in the N/Tert-1 lines used for this analysis, and assessed GO enrichment in relation to these [88]. Again, fibroblasts enhanced p53 and γH2AX protein expression in all N/Tert-1 lines, while pRb was enhanced in N/Tert-1 and N/Tert-1+E6E7 (Figure 5A). GO enrichment revealed that *TP53* was not enhanced at the RNA expression level, indicating that fibroblast enhancement of p53 protein expression, is likely mediated at the level of translation, post-translation, or protein stability, however, some p53 inducible proteins did appear to be regulated at the level of RNA (GO enrichment Figure 5B, p53 qPCR time course validation 5C-E) [88]. *TP53I13, TP53TG1*, and *TP53TG5* overexpression have been linked to the inhibition of cell proliferation and tumor suppression [143–145]. Enhancement of these tumor suppressors in N/Tert-1+HPV16 grown in the presence of J2, again suggests that fibroblasts promote a less transformed genotype (Figure 5B). GO enrichment related to Rb signaling is less clear. However, the observed *RB1, RBL1, RB1CC1, RBBP4P1* RNA upregulation (Figure 5F) in N/Tert-1+HPV16 grown in the presence of J2, is suggestive of a less transformed genotype [146–149]. *H2AX* RNA upregulation was demonstrated in both N/Tert-1+E6E7 and N/Tert-1+HPV16 grown in the presence of fibroblasts (Figure 5G), indicating a partial role in the previously observed J2 enhancement of γH2AX protein (the phosphorylated form of the H2AX variant) [88].

**Figure 5.**
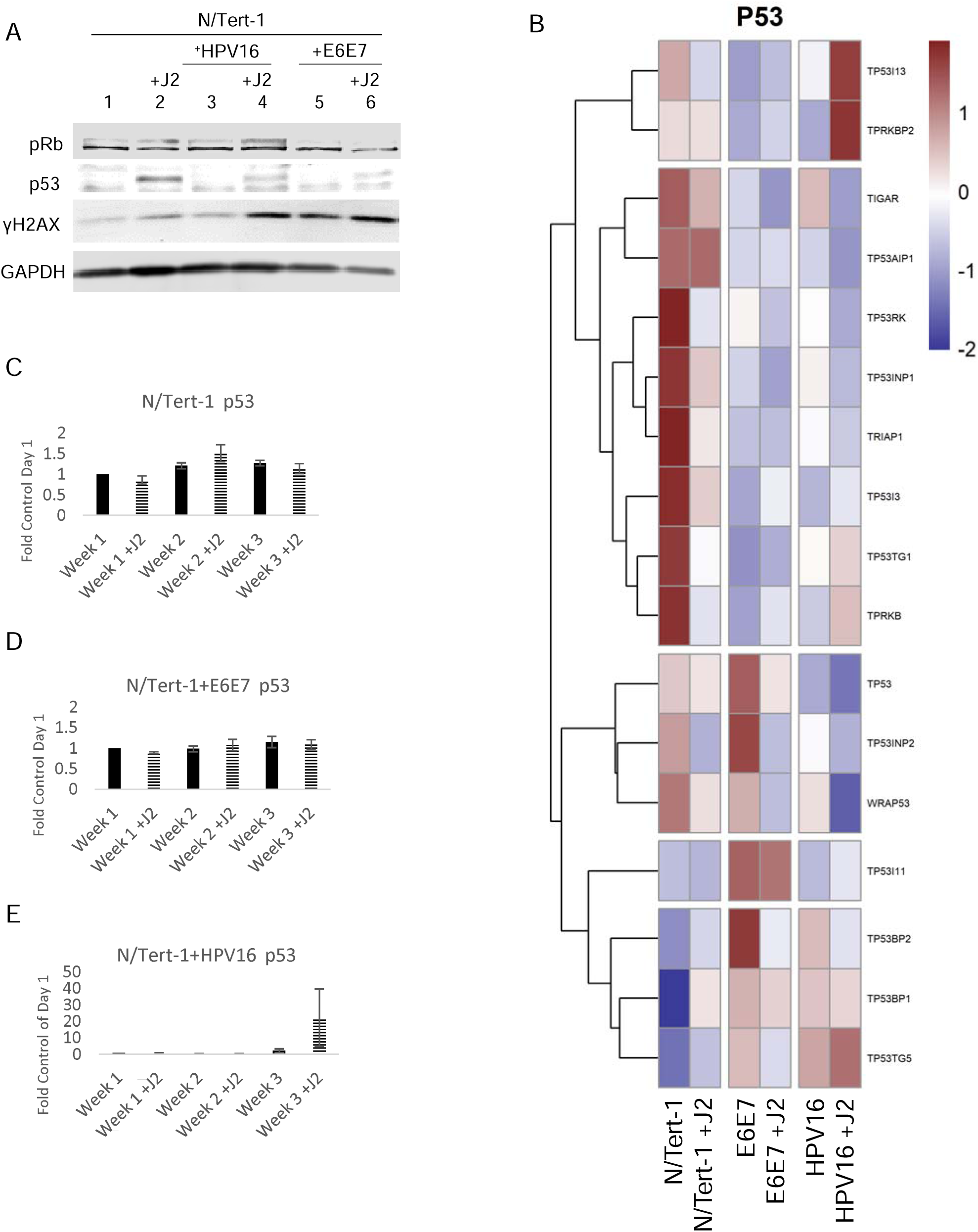
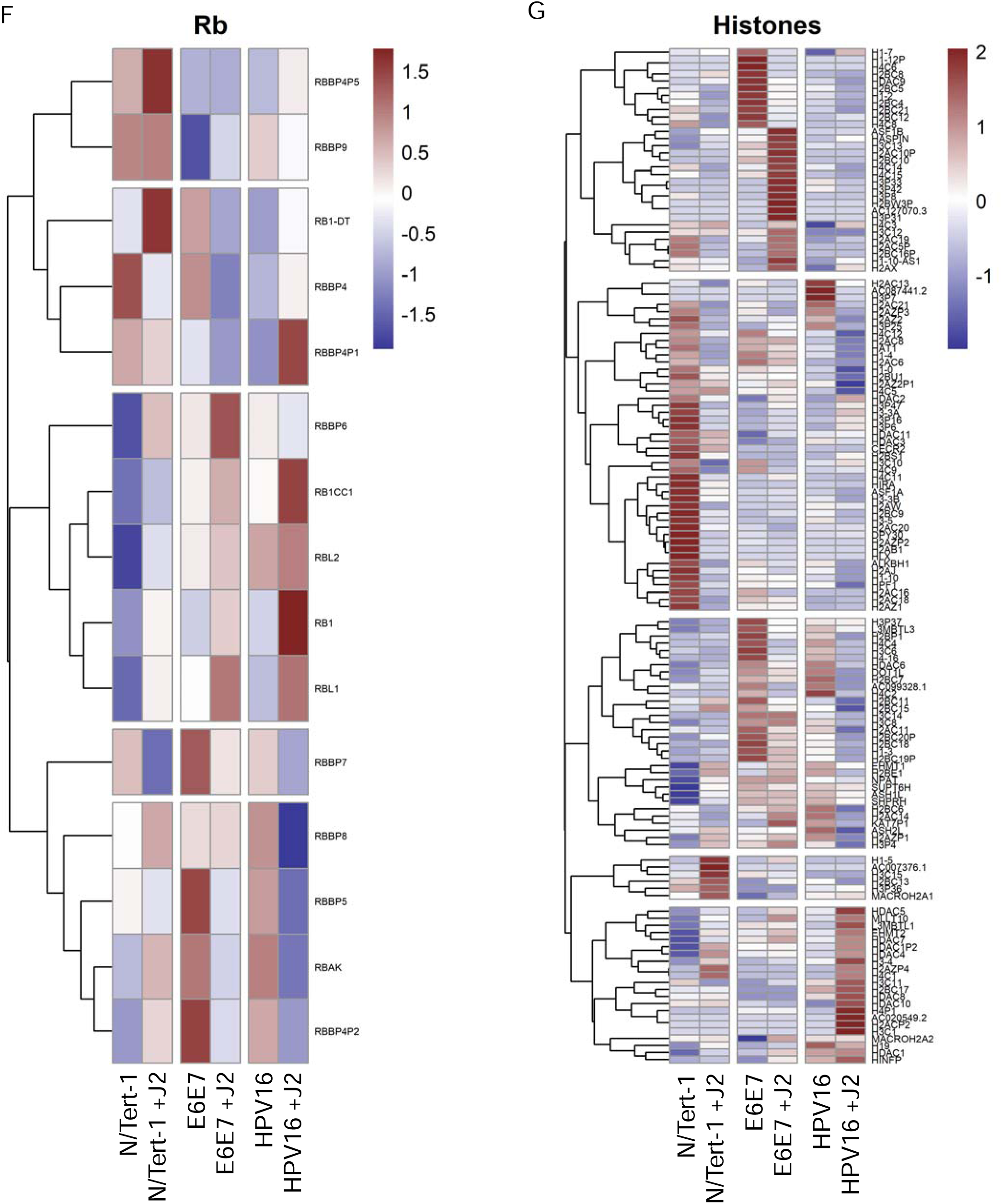
Fibroblasts differentially regulate p53, pRb, and histone related expression. **5A.** N/Tert-1 (lanes 1,2) N/Tert-1+E6E7 (lanes 3,4), N/Tert-1+HPV16 (lanes 5,6) cells were seeded on day 0 and grown in the presence or absence of J2s for 1 week. Cells were washed to remove J2s in noted conditions, trypsinized, lysed, and analyzed via western blotting for pRb, p53, and γH2AX. GAPDH was utilized as a loading control. **5B.** Heat map demonstrating significant p53 GO enrichment all groups. **5C.** N/Tert-1, **5D.** N/Tert-1+E6E7, and **5E.** N/Tert-1+HPV16 were grown in the presence or absence of J2s for 3 weeks. Time course of p53 RNA is presented at fold control of day 1. **5F.** Heat map demonstrating significant pRb RNA enrichment all groups. 5G. Heat map demonstrating significant histone RNA enrichment in all groups.

Another significant observation from our GO enrichment cross-comparisons were alterations in genes associated with cell cycle regulation and progression (Figure 6). Cell cycle regulation and progression are notably altered during oncogenic transformation and HPV-related transformation [1,150–152]. N/Tert-1+E6E7 cells cocultured with fibroblasts, markedly upregulated GO enrichment related to cell cycle regulation, cell cycle progression, cell division, and mitotic progression; these alterations were highly suggestive of significant transformation (Figures 6A-G)[153–155]. Conversely, N/Tert-1+HPV16 grown in the presence of J2 upregulated GO enrichment in tissue development that was highly suggestive of a less transformed genotype (Figure 6H). In particular, the expression of *KRT4* and *KRT13* decreases in transformed epithelial cells; N/Tert- 1+HPV16 grown in the presence of J2 instead showed enhanced *KRT13* and *KRT4* levels [156]. Likewise, HPV16+ keratinocytes maintained in J2 exhibited enhanced stress response GO enrichment, including the upregulation of a number of genes related to tumor suppression (Figure 6I). Again, highlighting the ability of fibroblasts to differentially regulated transformative genotypes.

**Figure 6.**
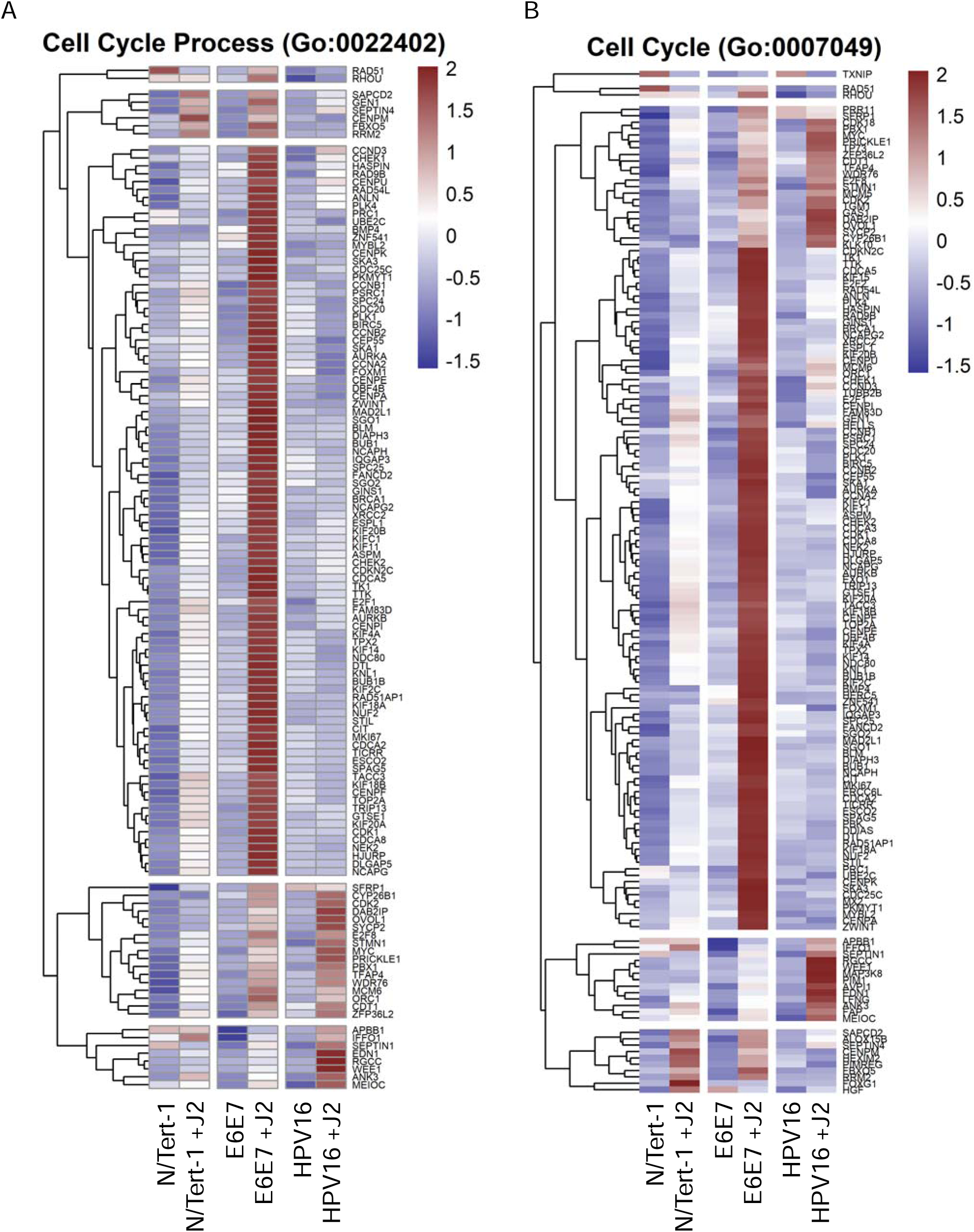
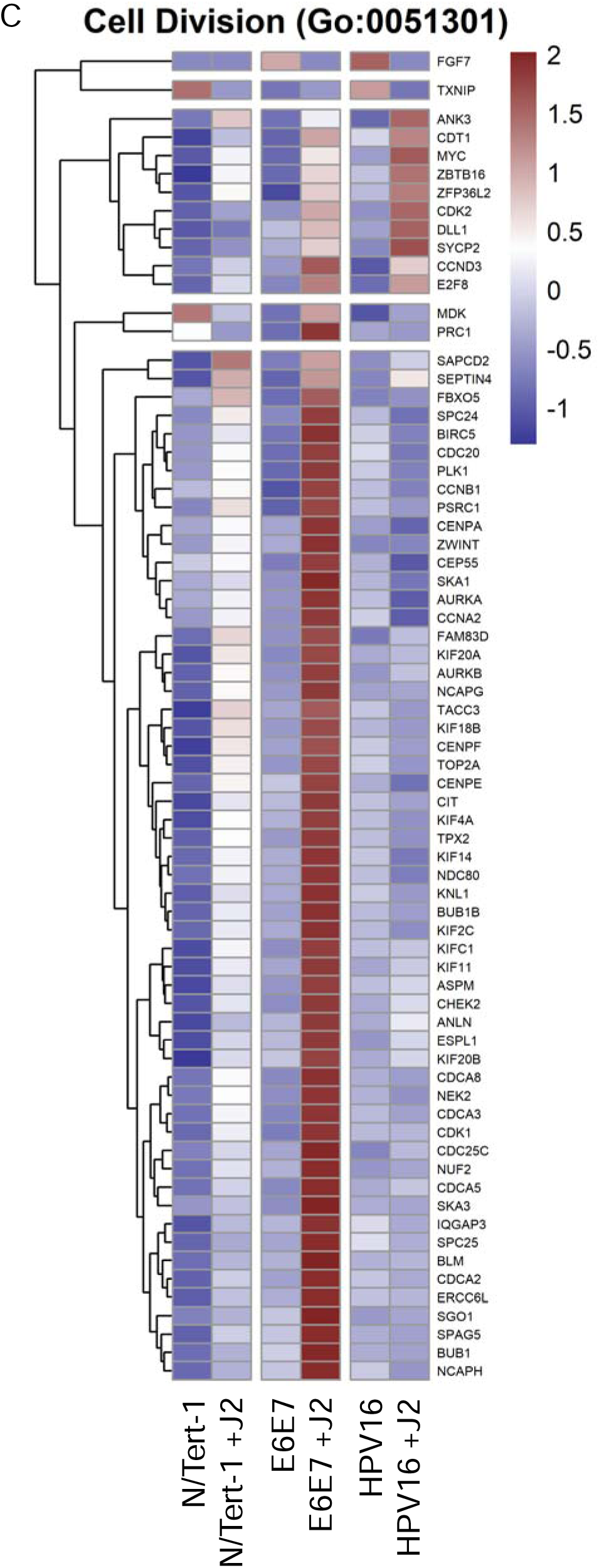
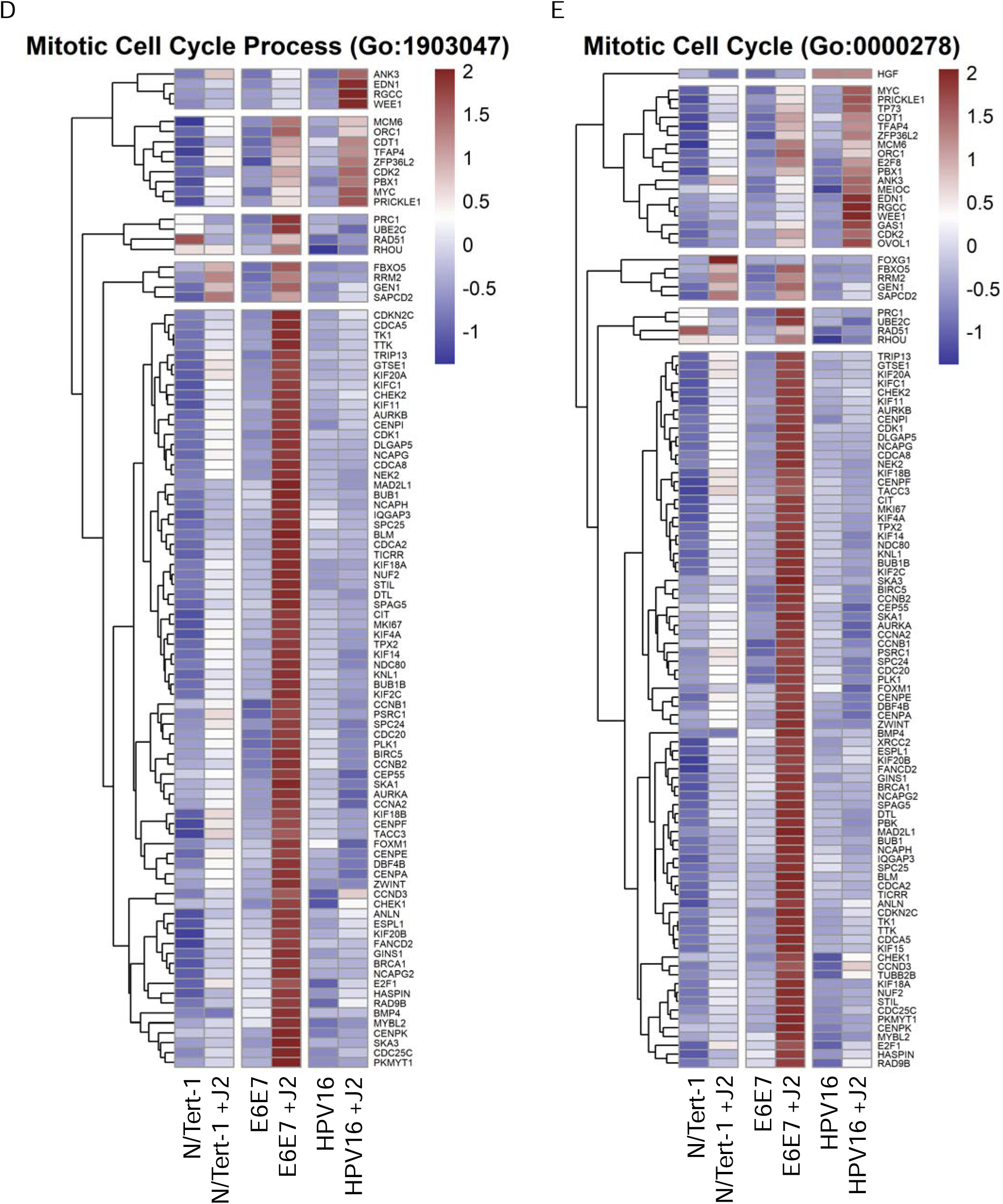
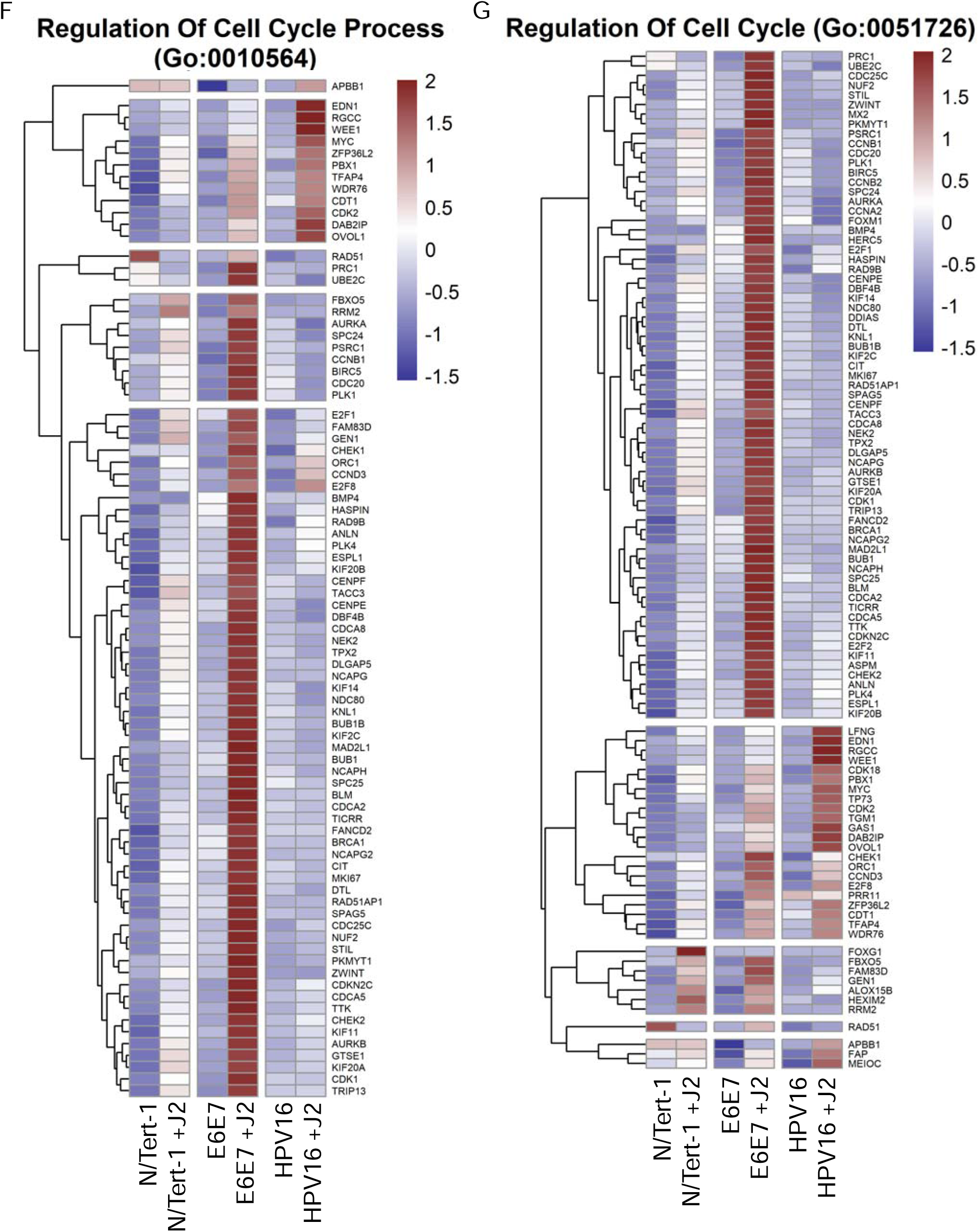
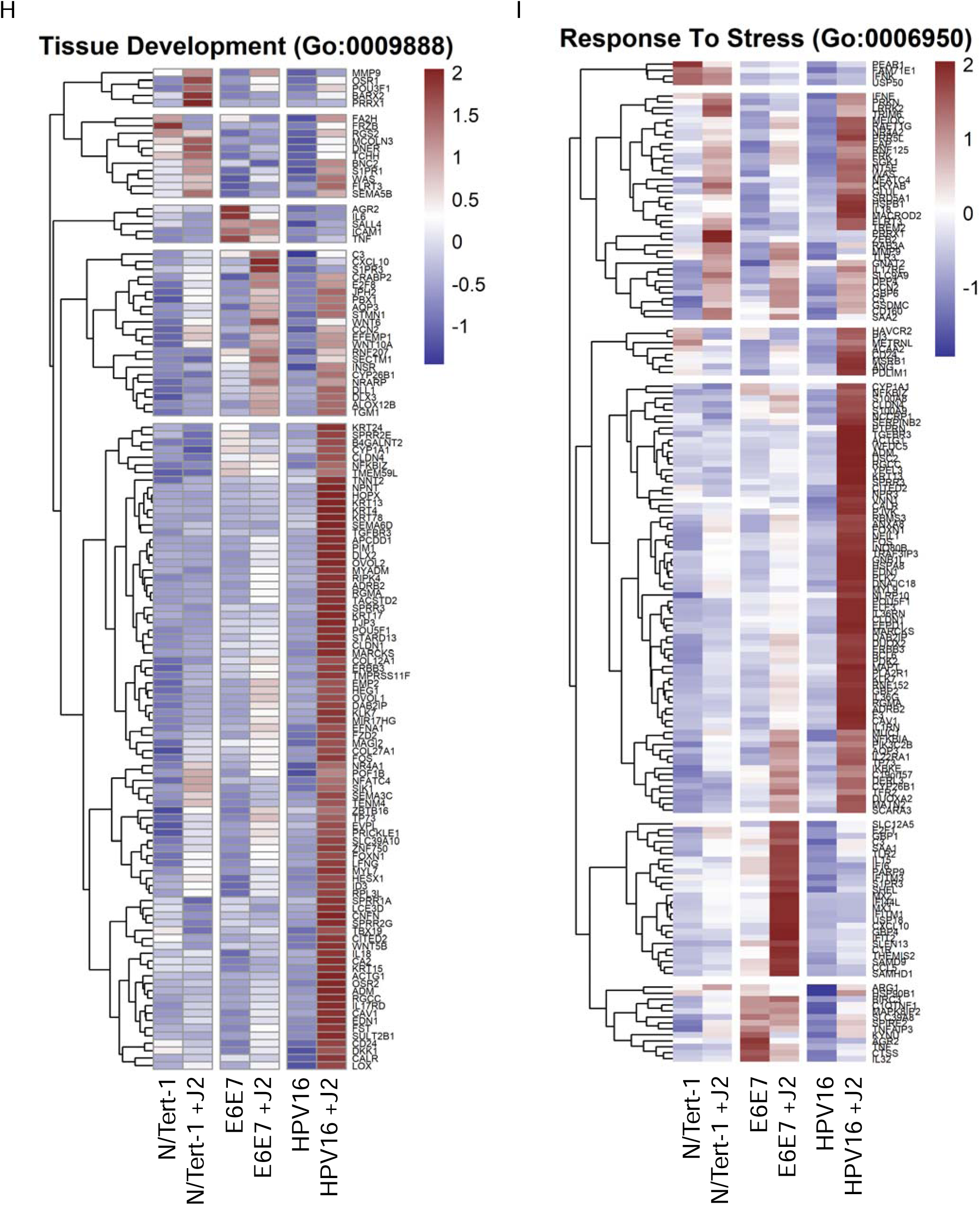
Fibroblasts differentially regulate cell cycle, tissue development, and stress response related GO enrichment. **6A.** Heat map demonstrating significant GO:0022402 cell cycle progression across all groups. **6B.** Heat map demonstrating significant GO:0007049 cell cycle across all groups. **6C.** Heat map demonstrating significant GO:0051301 cell division across all groups. **6D.** Heat map demonstrating significant GO:1903047 mitotic cell cycle progress across all groups. **6E.** Heat map demonstrating significant GO:0000278 mitotic cell cycle across all groups. **6F.** Heat map demonstrating significant GO:0010564 regulation of cell cycle process across all groups. **6G.** Heat map demonstrating significant GO:0051726 regulation of cell cycle across all groups. **6H.** Heat map demonstrating significant GO:0009888 tissue development across all groups. **6I.** Heat map demonstrating significant GO:0006950 response to stress across all groups.

### 3.2 Differential HPV RNA Expression Altered by Fibroblasts in Keratinocytes

We and others have demonstrated the importance of fibroblast co-culture for viral episome maintenance in HPV+ keratinocytes [87,88,122]. As previously demonstrated in HFK+HPV16, N/Tert-1+HPV16 grown in the presence of fibroblasts for one week demonstrated significantly enhanced integration events in the absence of J2 (Figure 7A) [88]. Mining of viral reads from RNA-seq data was performed and interpreted utilizing a technique previously developed [17,157,158]. RNA differential expression analysis demonstrated that N/Tert-1+HPV16 grown in the presence of J2 had significantly higher levels of *E2, E5, E6*, and *E7* transcripts than cells grown in the absence of J2 (RNA-seq reads in Figure 7B, *E2*, and *E6* qPCR time course validation in Figures 7C and 7D, respectively). Alternatively, N/Tert-1+E6E7 grown in the presence of J2 expressed lower RNA transcripts of *E7*, and similar *E6* transcripts in comparison to cells grown in the absence of J2 (RNA-seq reads in Figure 7B and *E6* qPCR time course validation in Figure 7E).

**Figure 7.**
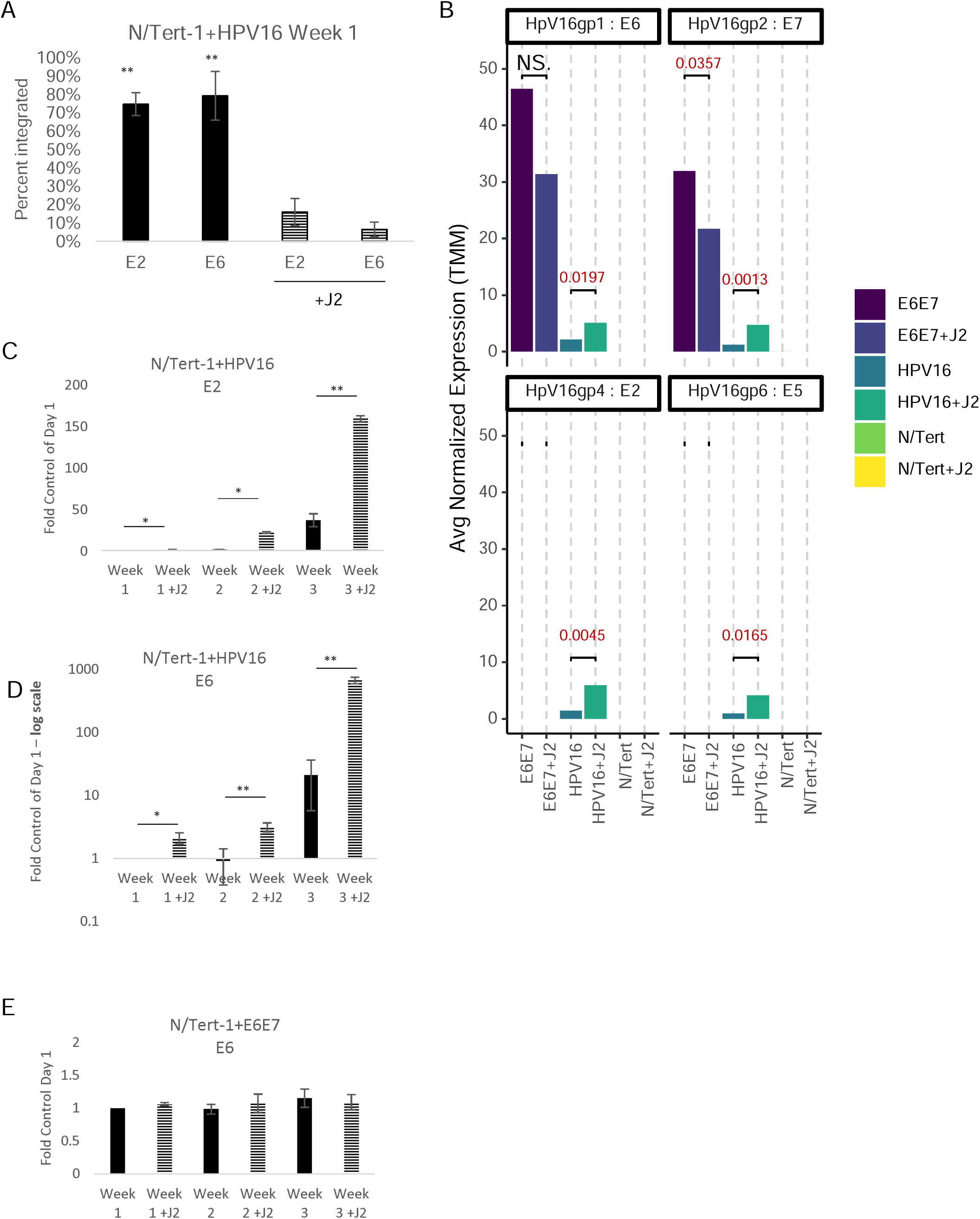
Fibroblasts support viral RNA expression and episomal maintenance in HPV+keratinocytes. **7A.** N/Tert-1+HPV16 cells were grown in the presence or absence of J2s for 1 week. Cells were washed to removed J2, then lysed and analyzed for DNA expression of E2 and E6 via the exonuclease V assay, in comparison to GAPDH and mitochondrial DNA controls. Results are presented as percent integration as calculated from the cut ratio of matched GAPDH. **P < 0.01. **7B.** Differential expression data from RNAseq from average normalized reads of E6, E7, E2, and E5 matched to HPV reference genome. Exact significance is presented for each (student’s t-test), NS represents no significance. **7C-E.** qPCR time course validation of E2 and E6 RNA expression in N/Tert-1+E6E7 and N/Tert-1+HPV16 in the presence or absence of J2 for 3 weeks, **7D** is presented in log scale. *P < 0.05. **P < 0.01.

### 3.3 Differential Proteomic Landscapes Altered by Fibroblasts in Keratinocytes

For label-free LC-MS/MS proteomic comparison, matched triplicate samples were harvested at the same time as RNA-seq; differential protein expression and bioinformatic analysis was performed, cross-matched to RNA-seq, and further assessed by bioinformatics. Processed datasets are available in Supplementary Data S4. Exact comparative analysis is presented as Venn diagrams in Figure 8 and comparative heatmaps in Figure 9. While mRNA expression precedes protein translation, the exact correlation between transcript levels and protein abundance is often poor; correlative assessments can instead be utilized for biomarker trends [159–162]. The Human Protein Atlas was first consulted to assess if comparative analysis supported our RNAseq observations that fibroblasts regulate the transformation potential in HPV+ keratinocytes [163–165]. Many oncogenic proteins were significantly downregulated in N/Tert- 1+HPV16 cells grown in the presence of J2; clinical pathology observations have proven that high expression of these proteins correlates with poor prognostics in either cervical cancer and/or head and neck cancer [163–165]. Fibroblast downregulation of these markers in N/Tert-1+HPV16 is suggestive of less transformation, which is in agreement with the observed changes in EMT markers in the RNA analysis. Global profiling of trends confirmed differentially regulated subgroups in relation to transformation events. Our overall observations suggest that fibroblasts influence genotypic profiles that support the viral lifecycle while inhibiting oncogenic progression in HPV+ keratinocytes. This fibroblast regulation pattern is inversed in E6E7+ keratinocytes, where oncogene expression is outside the control of E2.

**Figure 8.**
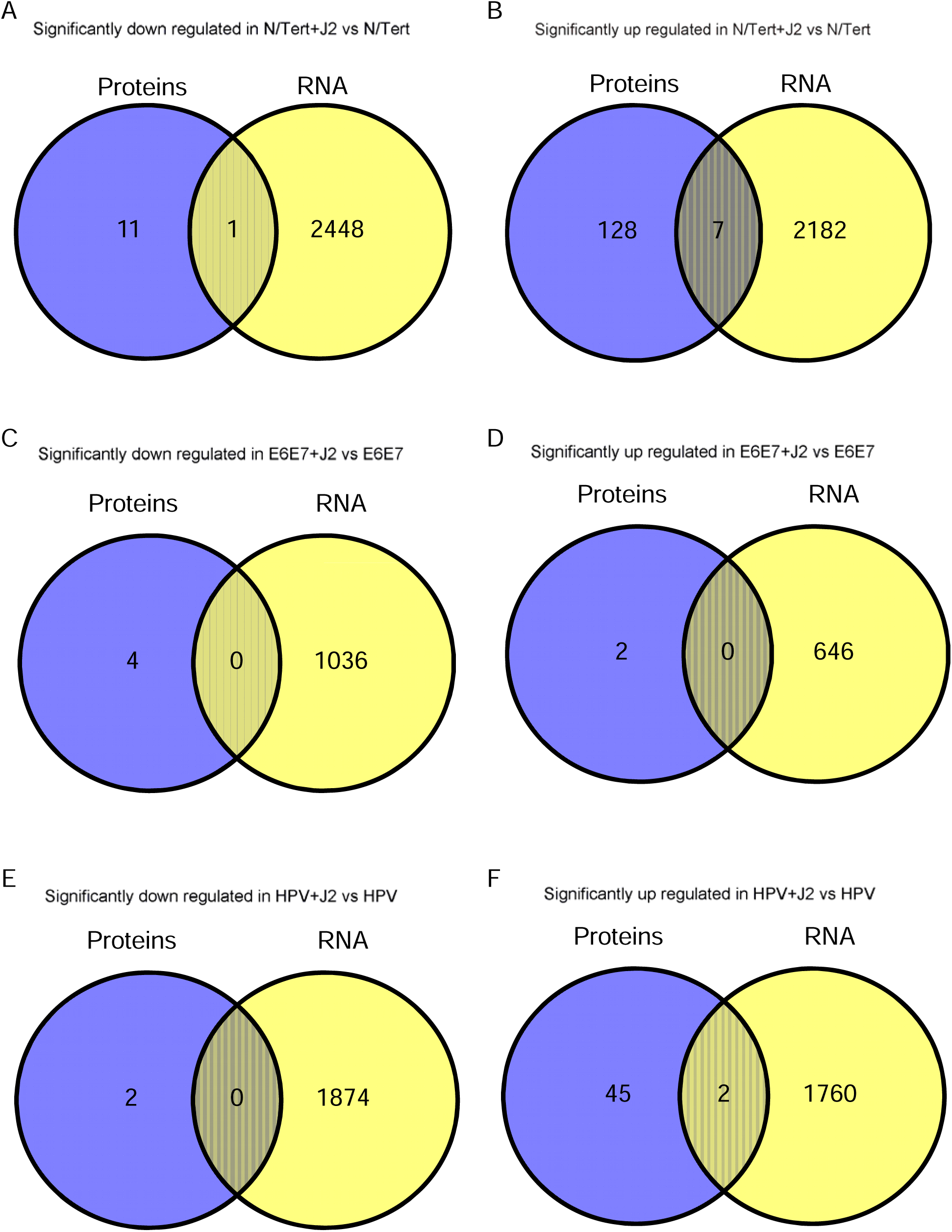
Differential expression Venn diagrams comparing significant up or down regulation via fibroblasts in RNA-seq and proteomic analysis. The sum of all the numbers in the circle represents the total number in the compared groups, and the overlapping area indicates the number of differential genes shared between the groups, as shown in the following figures. **8A,B.** Cross comparison of N/Tert-1 downregulation, and upregulation, respectively via fibroblasts. **8C,D.** Cross comparison of N/Tert-1+E6E7 downregulation, and upregulation, respectively via fibroblasts. **8E,F.** Cross comparison of N/Tert-1+HPV16 downregulation, and upregulation, respectively via fibroblasts.

**Figure 9.**
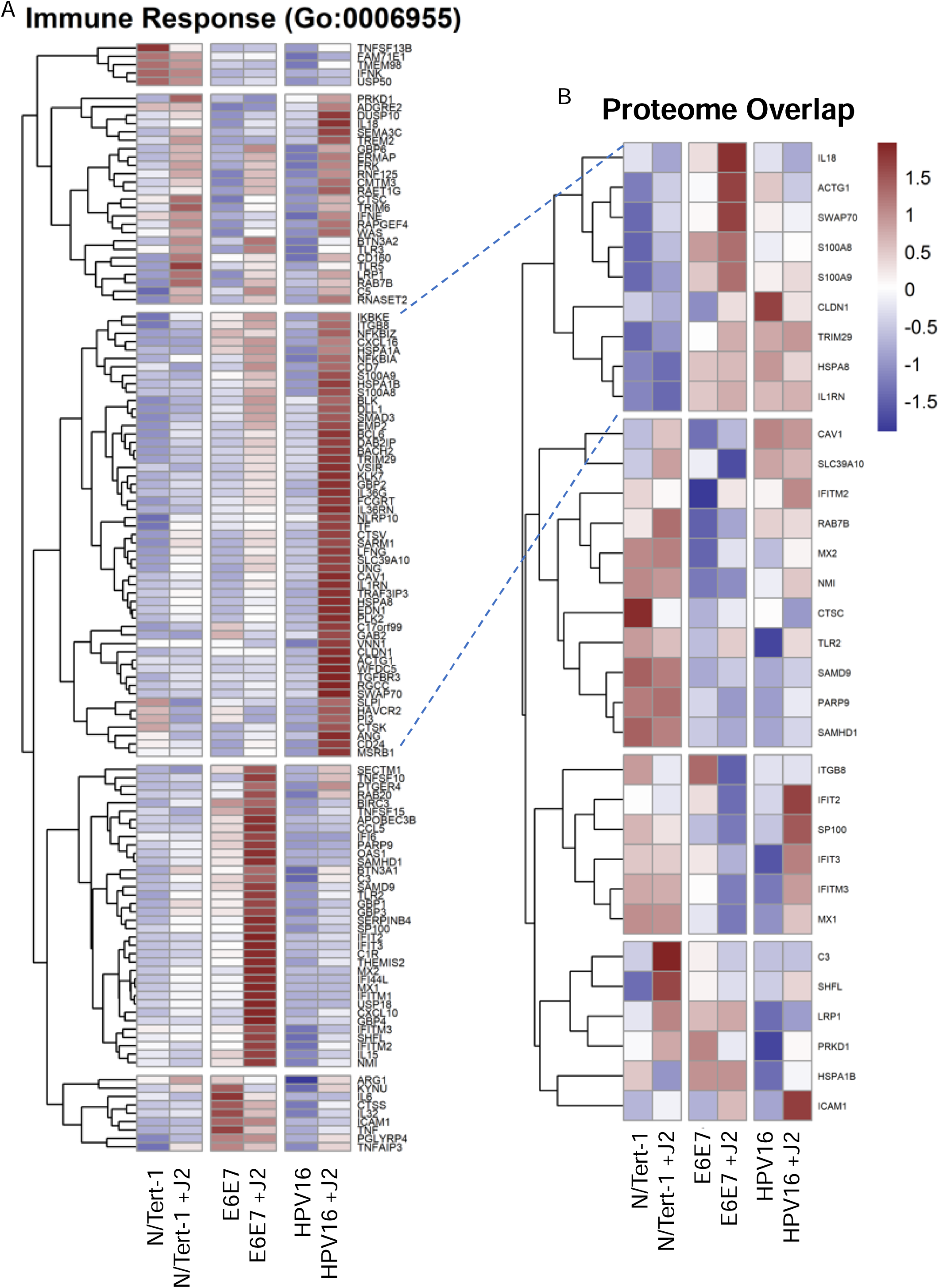
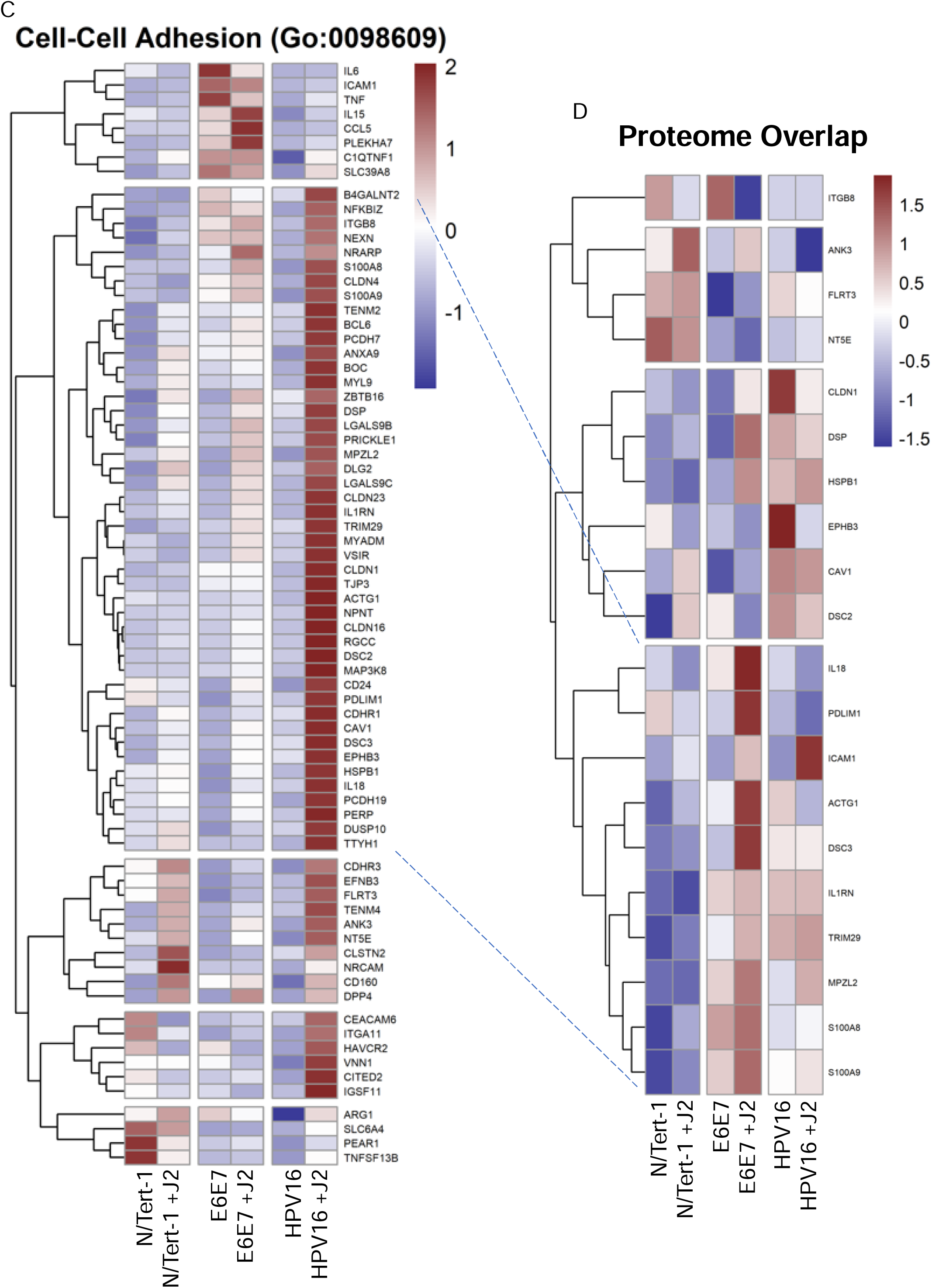
RNA-seq and proteomic cross comparisons demonstrate fibroblasts differentially regulate GO enrichment in relation to innate immune function and cell-cell adhesion. **9A.** Heat map demonstrating significant GO:0006955 immune response across all groups. **9B.** Matched heat map analysis of significant proteome alterations of GO:0006955 across all groups. **9C.** Heat map demonstrating significant GO:0098609 cell-cell adhesion across all groups. 9D. Matched heat map analysis of significant proteome alterations of GO:0098609 across all groups. Dotted lines are added to help visually compare similar matched sets.

## 4 Discussion

Decades of research have continued to improve the model systems utilized to mimic HPV infection and progression. Despite the increasing availability of improved models, a current challenge in the field is that these disease models still do not fully replicate the tissue complexity of the various epithelial sites where severe diseases develop [24,166]. The addition of fibroblast feeder cells for the generation of epithelial cell lines has improved both the efficiency of immortalization attempts, as well as contributing to tissue complexity in 2D growth settings [69,70]. Primary keratinocyte lines are easily generated for many epithelial sites of HPV infections, however primary cell lines do not allow for longitudinal studies [167]. Primary cultures can be immortalized with HPV; however, “control” cell lines are limited due to the nature of primary cell culture. Immortalized primary human keratinocytes using the catalytic subunit of telomerase (hTERT) have been generated for use as longitudinal “control” cell lines, however expression of hTERT alone is often insufficient for the immortalization of human keratinocytes [168]. Successfully immortalized keratinocyte lines like telomerase (hTERT) immortalized primary foreskin keratinocytes (N/Tert-1), the spontaneously immortalized normal immortal keratinocytes (NIKS), or the adult epidermis cell line generated from the periphery of a malignant melanoma (HaCaT) are thus utilized as surrogates for long term “control” comparisons [168,169]. HPV E6 and E7 can likewise be exploited to immortalize keratinocytes with improved efficiency, however, they are no longer completely null of HPV [57,71,170]. To assess how fibroblasts modulate viral-keratinocyte interactions, we carefully evaluated the most effective approach to control for all relevant factors. For this reason, we chose to utilize our well-characterized and matched N/Tert-1 keratinocyte lines [28,29,74,88,95,171].

Genomic and proteomic assessments in short-term 2D cultures revealed that fibroblasts promoted a less transformed state in N/Tert-1+HPV, whereas N/Tert-1+E6E7 may be more transformed in the presence of fibroblasts. The exact nature of oncogenic transformation remains largely speculative, although a number of biomarkers are well characterized in this progression [46,57,60,66,77,122,129,131,170,172]. Our studies confirmed that N/Tert-1+HPV maintained in fibroblasts sustained HPV episomes, consistent with a less progressed HPV genotypic state (Figure 7A) [77,79,80,88,158]. Likewise, host expression of host signaling regulation, was also suggestive of a less transformed state; specifically tight junction regulation, CXC chemokine expression, TNF-related signaling, and *TWIST* expression were most compelling (Figure 4). Conversely, when comparing the signaling regulation of N/Tert-1+E6E7 maintained in fibroblasts, the genotypic regulation presented the biological antithesis of the aforementioned observations (Figure 4). Additionally, N/Tert-1+E6E7 maintained in fibroblasts exhibited significant enhancement of cell cycle regulation that was suggestive of transformation (Figure 6). True longitudinal HPV transformation has yet to be demonstrated in traditional cell culture; our observations suggest that alterations in cell culture maintenance conditions are worth consideration for future analysis.

Organotypic raft cultures have also been used for the broad examination of how high-risk HPVs may drive neoplasia and cancer [166]. It is well noted that fibroblasts serve a fundamental role in epithelial differentiation and the viral lifecycle in this 3D model [23,43,44,173–175]. While 3D cultures present a model for reconstructing the viral lifecycle, these cultures are not useful for traditional cell maintenance. Likewise, 2D culture models can also be utilized to examine the viral lifecycle employing a calcium gradient medium, but differentiation also presents finite time points [166]. Future studies in our lab will extrapolate the transformation-related alterations presented, and assess how fibroblasts continue to regulate viral-host interactions temporally, spatially, and in the context of differentiation. These alterations will be considered at various stages of transformation, in 2D and 3D models, and in the context of both normal and cancer-associated fibroblasts.

## 5 Conclusion

Both our research and that of others have shown that interactions between fibroblasts and keratinocytes in HPV models are critical for maintaining episomal HPV genomes, influencing keratinocyte differentiation, and regulating viral transcription [23,43,44,52,88,121,173–175]. Here we present RNAseq analysis revealing that fibroblasts may regulate the transformation potential in HPV+ keratinocytes by regulating cytokine activity, cell junction proteins, and innate immune signaling. Proteomic analysis further supported these findings, highlighting fibroblasts’ ability to modulate protein expression linked to oncogenic transformation. Overall, fibroblasts were found to influence both viral and host cell signaling, promoting HPV lifecycle maintenance while potentially limiting cancer progression in HPV+ keratinocytes; conversely, E6E7+ keratinocytes were more transformed in the presence of fibroblasts and may present a more neoplastic model.

## Data availability statement

Following the 2023 NIH data management and sharing policy, all data resulting from the development of projects will be available in scientific communications presented at conferences and in manuscripts that will be published in peer-reviewed scientific journals. Data will be deposited in the Open Science Framework (OSF) platform. OSF can be accessed at https://osf.io. VCU is an OSF institutional member, and OSF is an approved generalist repository for the 2023 NIH data management and sharing policy.

## CRediT authorship contribution statement

**Claire D. James**: Writing – review & editing, Writing – original draft, Supervision, Methodology, Investigation, Formal analysis, Data curation. **Rachel L. Lewis**: Methodology, Investigation, Data curation, Validation. **Austin J. Witt**: Methodology, Investigation, Data curation, Validation. **Christiane Carter**: Writing – review & editing, Software, Methodology, Investigation, Formal analysis, Validation. **Nabiha M. Rais**: Methodology, Investigation, Data curation. **Xu Wang:** Formal analysis, Data curation. **Molly L. Bristol**: Writing – review & editing, Writing – original draft, Supervision, Resources, Project administration, Methodology, Investigation, Data curation, Funding acquisition, Formal analysis, Conceptualization, Validation, Visualization.

## Acknowledgments

This work was supported by the VCU Philips Institute for Oral Health Research, the VCU Quest Fund, the National Institute of Dental and Craniofacial Research/NIH/DHHS R03 DE029548, and the National Cancer Institute-designated Massey Cancer Center grant P30 CA016059. Services in support of the research project were provided by the VCU Massey Comprehensive Cancer Center Bioinformatics Shared Resource. Massey is supported, in part, with funding from NIH-NCI Cancer Center Support Grant P30 CA016059. Services and products in support of the research project were generated by the VCU Massey Comprehensive Cancer Center Proteomics Shared Resource, supported, in part, with funding from NIH-NCI Cancer Center Support Grant P30 CA016059.

